# Cytomegaloviral determinants of CD8^+^ T cell programming and RhCMV/SIV vaccine efficacy

**DOI:** 10.1101/2020.09.30.321349

**Authors:** Daniel Malouli, Scott G. Hansen, Meaghan H. Hancock, Colette M. Hughes, Julia C. Ford, Roxanne M. Gilbride, Abigail B. Ventura, David Morrow, Kurt T. Randall, Husam Taher, Luke S. Uebelhoer, Matthew R. McArdle, Courtney R. Papen, Renee Espinosa Trethewy, Kelli Oswald, Rebecca Shoemaker, Brian Berkemeier, William J. Bosche, Michael Hull, Justin M. Greene, Michael K. Axthelm, Jason Shao, Paul T. Edlefsen, Finn Grey, Jay A. Nelson, Jeffrey D. Lifson, Daniel Streblow, Jonah B. Sacha, Klaus Früh, Louis J. Picker

**Affiliations:** Vaccine and Gene Therapy Institute and Oregon National Primate Research Center, Oregon Health & Science University, Beaverton, OR 97006; AIDS and Cancer Virus Program, SAIC Frederick, Inc., Frederick National Laboratory, Frederick, MD 21702; Population Sciences and Computational Biology Programs, Fred Hutchinson Cancer Research Center, Seattle, WA 98109; Division of Infection and Immunity, Roslin Institute, The University of Edinburgh, Edinburgh, United Kingdom

## Abstract

Simian immunodeficiency virus (SIV) insert-expressing, 68-1 Rhesus Cytomegalovirus (RhCMV/SIV) vectors elicit major histocompatibility complex (MHC)-E- and -II-restricted, SIV-specific CD8^+^ T cell responses, but the basis of these unconventional responses and their contribution to demonstrated vaccine efficacy against SIV challenge in the rhesus monkeys (RMs) has not been characterized. We demonstrate that these unconventional responses resulted from a chance genetic rearrangement in 68-1 RhCMV that abrogated the function of eight distinct immunomodulatory gene products encoded in two RhCMV genomic regions (Rh157.5/.4 and Rh158-161). Differential repair of these genes with either RhCMV-derived or orthologous human CMV (HCMV)-derived sequences (UL128/130; UL146/147) leads to either of two distinct CD8^+^ T cell response types – MHC-Ia-restricted-only, or a mix of MHC-II- and MHC-Ia-restricted CD8^+^ T cells. Despite response magnitude and functional differentiation being similar to RhCMV 68-1, neither alternative response type mediated protection against SIV challenge. These findings implicate MHC-E-restricted CD8^+^ T cell responses as mediators of anti-SIV efficacy and indicate that translation of RhCMV/SIV vector efficacy to humans will likely require deletion of all the genes that inhibit these responses from the HCMV/HIV vector.

**One-sentence summary:** Eight genes in two spatially distinct RhCMV gene regions control induction of unconventionally restricted CD8^+^ T cell responses and the efficacy of RhCMV/SIV vaccine vectors against SIV challenge.

---

Among viruses, CMVs are unique in their ability to elicit and indefinitely maintain high frequency CD4^+^ and CD8^+^ T cell responses with broad tissue distribution and prominent effector memory differentiation (EM-bias) (*1–3*). CMV-based vaccine vectors were developed to exploit this immunobiology, addressing the hypothesis that such circulating and tissue-based, effector-differentiated T cell responses, directed at heterologous pathogens via vaccination, would provide for a rapid immune effector intercept of nascent infections, prior to effective pathogen immune evasion (*4, 5*). This concept has been validated in the SIV-RM model of AIDS, where, across multiple studies, 50-60% of RMs vaccinated with RhCMV/SIV vectors show early stringent control (replication arrest) of highly pathogenic SIVmac239, with the vast majority of protected RM going on to eventually clear the infection (*6–8*). Although this “control and clear” pattern of anti-SIV efficacy, which has not been observed with any other vaccine modality, is consistent with an early CD8^+^ T cell effector intercept of infection by EM CD8^+^ T cells, the unexpected finding that, in contrast to natural RhCMV infection, the epitopes targeted by 68-1-based RhCMV vector-elicited CD8^+^ T cells are restricted by MHC-II and MHC-E, and not classical polymorphic MHC-Ia, and include universal epitopes (“supertopes”) of both restriction types (*9, 10*), has suggested that an early pathogen intercept may not be the only unique mechanism underling the efficacy of these vectors. The 68-1 RhCMV strain was extensively passaged on fibroblasts prior to cloning the viral genome as a bacterial artificial chromosome (BAC) for vector construction, and similar to fibroblast-adapted HCMV strains, had developed genetic mutations that inactivated the pentameric receptor complex (PRC), a CMV entry receptor for non-fibroblast cell types that is selected against during fibroblast passage (*11, 12*). In the case of 68-1 RhCMV, PRC inactivation occurred via deletion of the genes Rh157.5 and Rh157.4, RhCMV orthologs of HCMV UL128 and UL130, due to a deletion in the flanking region of an inverted genome segment (*11*) (**Fig. 1A**). Repair of Rh157.5 and Rh157.4 gene expression in clone 68-1.2 of RhCMV (*13*) restored conventional MHC-Ia-restricted-only CD8^+^ T cell recognition (*10*) (**Fig. 1B**), suggesting a role of the PRC in this unique reprogramming of vector immunogenicity. Here, we sought to further characterize the genetic and mechanistic basis of 68-1 RhCMV’s unconventional CD8^+^ T cell response programing, in particular the role of the PRC, and determine whether unconventional MHC restriction is necessary for vaccine efficacy, with the ultimate goal of defining the vector genotype necessary for clinical development of an efficacy-optimized HCMV-vectored HIV vaccine.

**Figure 1.**
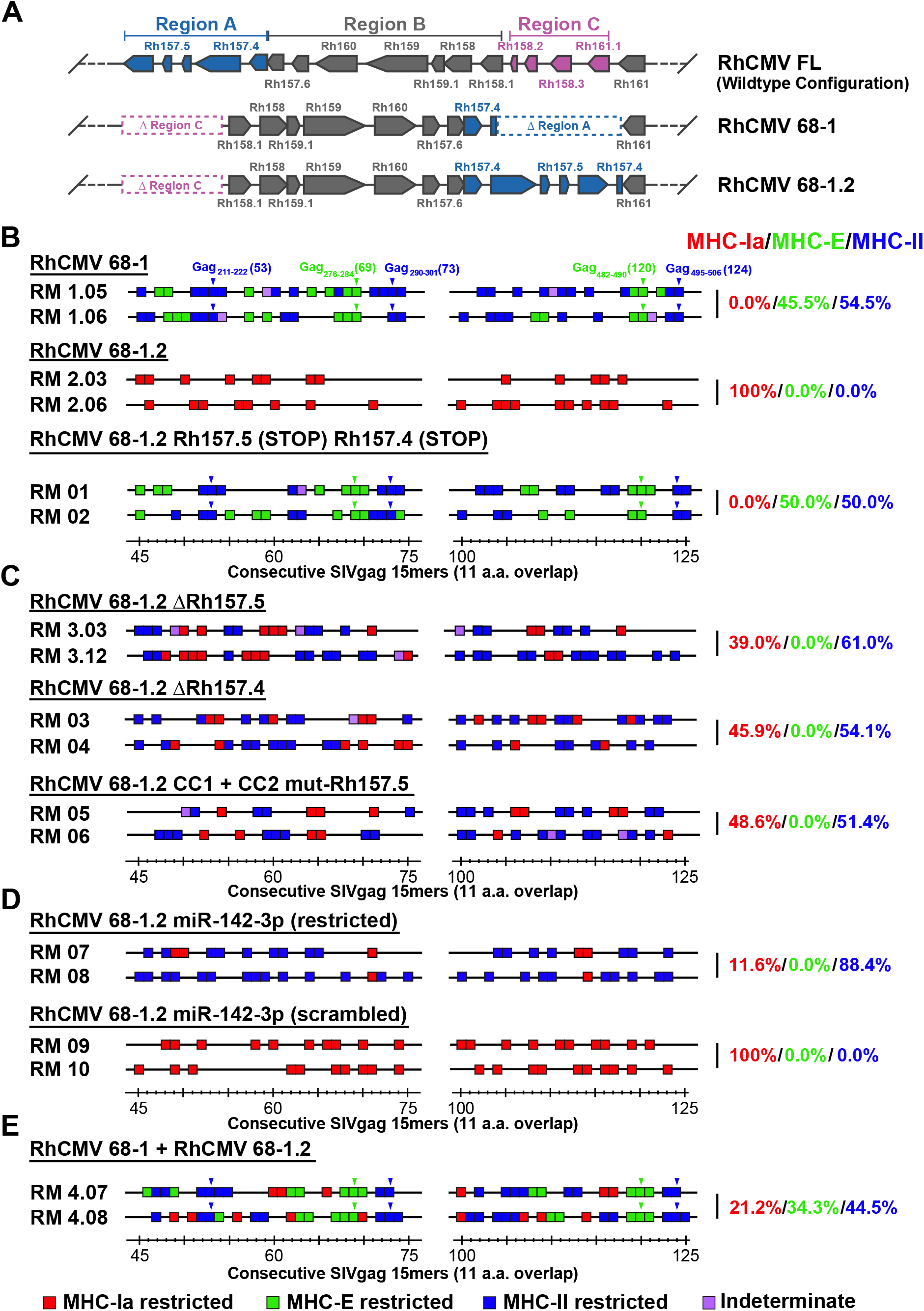
Role of the PRC in RhCMV vector CD8^+^ T cell response programming. **(A)** Schematic of the genetic differences between 68-1, 68-1.2, and FL (wildtype) RhCMV in the region of the genome encoding the PRC, showing the inversion-deletion event that occurred during *in vitro* passage of the 68-1 strain and its partial repair in 68-1.2. Diagrams of all modified versions of 68-1, 68-1.2 and FL RhCMV vectors are shown in **fig. S1** (note that all versions were engineered to express SIVgag). **(B-E)** MHC restriction analysis of SIVgag-specific CD8^+^ T cell responses elicited by the designated RhCMV vectors. Peripheral blood CD8^+^ T cells from 2-15 RM inoculated with each of these vectors were assessed by a flow cytometric intracellular cytokine staining (**ICS**) assay (based on induction of TNF and/or IFN-γ) for reactivity with each of 125 consecutive 15mer peptides (with 11 amino acid overlap) comprising the SIVgag protein sequence, with all above-threshold single 15mer responses depicted by boxes in the figure. Each response was then further analyzed for MHC-restriction by blocking separately with anti-MHC-I mAb W6/32, the MHC-E blocking peptide VL9, and the anti-MHC-II mAb G46-6. Assignment of MHC-Ia-, MHC-E-, and MHC-II-restriction of each response was based on response blocking by W6/32 alone (red boxes), W6/32 and VL9 alone (green boxes), and G46-6 alone (blue boxes), respectively, with responses not meeting these criteria labeled indeterminate (purple) (*9, 10*). In addition, CD8^+^ T cells from all RM were evaluated for responsiveness to the MHC-E supertope peptides SIV Gag_276-284_ (15mer #69) and Gag_482-490_ (#120) and the MHC-II supertope peptides SIV Gag_211-222_ (#53), Gag_290-301_ (#73), and Gag_495-506_ (#124), with above threshold responses to these supertopes indicated by green and blue arrowheads, respectively. The figure shows only the SIVgag regions from amino acid 45-75 and amino acid 100-125 (the regions including the supertopes) of representative RM (for those vectors studied with >2 RM), but complete epitope analysis results are presented in **table S1**. These overall data were used to calculate the % of the total MHC restriction-assignable SIVgag epitopes that were MHC-Ia-, MHC-E-, and MHC-II-restricted (shown at right in red/green/blue, respectively).

The PRC is a CMV entry receptor for endothelial, epithelial, and of particular relevance for T cell priming, myeloid-derived cells such as macrophages and dendritic cells, and PRC abrogation reduces infection of these cell types (*12*). To determine the role of the PRC and tropism in preventing 68-1.2 RhCMV vectors from eliciting unconventional responses, we constructed and tested a series of PRC- and non-PRC, tropism-modified 68-1.2 RhCMV/gag vectors, asking whether these modifications resulted in changes in the epitope targeting of vaccine-induced SIVgag-specific CD8^+^ T cell responses (**figs. S1,2; Table S1**). First, we demonstrated that incorporation of stop mutations into the Rh157.5 and Rh157.4 genes converted CD8^+^ T cell responses from targeting MHC-Ia-restricted epitopes (wildtype and 68-1.2 immunotype) to MHC-II- and MHC-E-restricted epitopes (68-1 immunotype), which demonstrates that protein expression, not alteration of genomic configuration by 68-1 inversion-deletion, is responsible for response programming (**Fig. 1B**). We then tested 68-1.2 vectors with genetic modifications that affect PRC function without complete abrogation of both Rh157.5 or Rh157.4 expression, including deletion (Δ) of Rh157.5 alone, Rh157.4 alone, and functional inactivation of the PRC by alanine-substitution mutation of two charged cluster (CC) regions of the UL128 ortholog Rh157.5 (*14*) (**figs. S1,2**). Surprisingly, the CD8^+^ T cell responses elicited by these 3 vectors did not recapitulate either the 68-1 or 68-1.2 CD8^+^ T cell response types, but instead resulted in a different response type characterized by mixed CD8^+^ T cell targeting of both MHC-Ia- and MHC-II-restricted epitopes, the latter notably without supertopes (**Fig. 1C**).

To determine whether this new response type was due to alteration of vector tropism, we engineered a PRC-independent tropism restriction into the 68-1.2 vector by incorporating target sites for the myeloid-specific microRNA (miR)-142-3p into the 3’ untranslated region (UTR) of essential viral genes (*15, 16*), which, by suppressing expression of essential viral genes, impedes the vector’s ability to productively infect miR-142-3p-expressing myeloid cells (e.g., simulating one possible effect of PRC inactivation relevant to CD8^+^ T cell response priming: decreased infection efficiency of myeloid-derived antigen presenting cells) (**fig. S3**). Notably, this miR-142-3p-restricted 68-1.2 vector elicited a very similar pattern of CD8^+^ T cell epitope targeting (MHC-Ia-+ MHC-II-restriction, without supertopes) as the ΔRh157.5, ΔRh157.4, and CC1+CC2 mutant-Rh157.5 68-1.2 vectors, except that the MHC-II-restricted CD8^+^ T cell priming of the former vector was more efficient (88% of total epitopes) than the latter vectors (57% of total epitopes; p<0.001 by binomial exact test)(**Figs. 1C,D**). Taken together, these data indicate that PRC inactivation alone is not responsible for the full development of the exclusively unconventionally targeted CD8^+^ T cell responses manifested by 68-1 vectors, but does provide for elicitation of non-supertope, MHC-II-restricted CD8^+^ T cells, an effect that is likely attributable to reduced infection of myeloid-derived cells driven by PRC inactivation. Additionally, these data indicate that both the Rh157.5 and Rh157.4 gene products have a PRC-independent activity that interferes with elicitation of CD8^+^ T cell responses to all MHC-E-restricted epitopes and to MHC-II-restricted supertopes, potentially related to their N-terminal homology with host chemokines (**fig S4**). This inhibitory activity operates at the level of the vector-infected cells, not systemically, as coadministration of 68-1 and 68-1.2 RhCMV/SIVgag vectors to RMs results in CD8^+^ T cell responses targeting epitopes of all three MHC restriction types: MHC-Ia, MHC-II and MHC-E, including supertopes (**Fig. 1E**).

We next asked the question of whether abrogating both the PRC and non-PRC-related immune programming activities of Rh157.5 and Rh157.4 was sufficient to convert a wildtype-like RhCMV, with conventional MHC-Ia-restricted CD8^+^ T cell targeting, to a 68-1-like vector with exclusively MHC-II- and MHC-E-restricted CD8^+^ T cell responses. Using sequence information from primary RhCMV isolates, including the original 68-1 isolate (*17*), we reconstructed a full length RhCMV clone (RhCMV FL) in which all suspected deletions, inversions, frameshifts and premature termination codons affecting viral open reading frames were repaired and the Rh13.1 open reading frame (ORF) was replaced by an SIVgag insert (*18*). RhCMV FL has robust *in vivo* spread (*18*), and as expected for a wildtype genetic configuration (*9*), elicits MHC-Ia-restricted, CD8^+^ T cell responses (**Fig. 2A**). To determine whether inactivation of Rh157.5 and Rh157.4 was sufficient for unconventional CD8^+^ T cell response programming, we deleted both genes from RhCMV FL. Unexpectedly, the resulting recombinant vector (RhCMV FL ΔRh157.5/Rh157.4) elicited CD8^+^ T cell responses showing the same conventional MHC-Ia-restriction as the parent RhCMV FL vector, suggesting that expression of genes outside the Rh157 region additionally interfere with the induction of unconventionally restricted CD8^+^ T cell responses (**Fig. 2A**). Since the inversion/deletion event in 68-1 also deleted genes orthologous to HCMV UL146 and UL147, which encode viral CXC chemokine-like proteins with similarity to CXCL-1 (*11*) (**Fig. 1A; fig. S4**), we constructed RhCMV FL vectors with deletions of the UL146/147 orthologs Rh158-Rh161 either alone or together with deletion of Rh157.5 and Rh157.4 (**fig. S1**). Deletion of the Rh158-Rh161 region alone (RhCMV FL ΔRh158-161) also did not change the MHC-Ia restriction of RhCMV FL elicited CD8^+^ T cell responses; however, deletion of *both* Rh157.5/Rh157.4 and Rh158-Rh161 [double-deleted (dd) RhCMV FL] did result in the same MHC-II-+ MHC-E-restricted CD8^+^ T cell targeting as the original 68-1 RhCMV vector (**Figs. 2A**), demonstrating that deletion of both these distinct regions from a typical circulating RhCMV strain is required for unconventional response programming.

**Figure 2.**
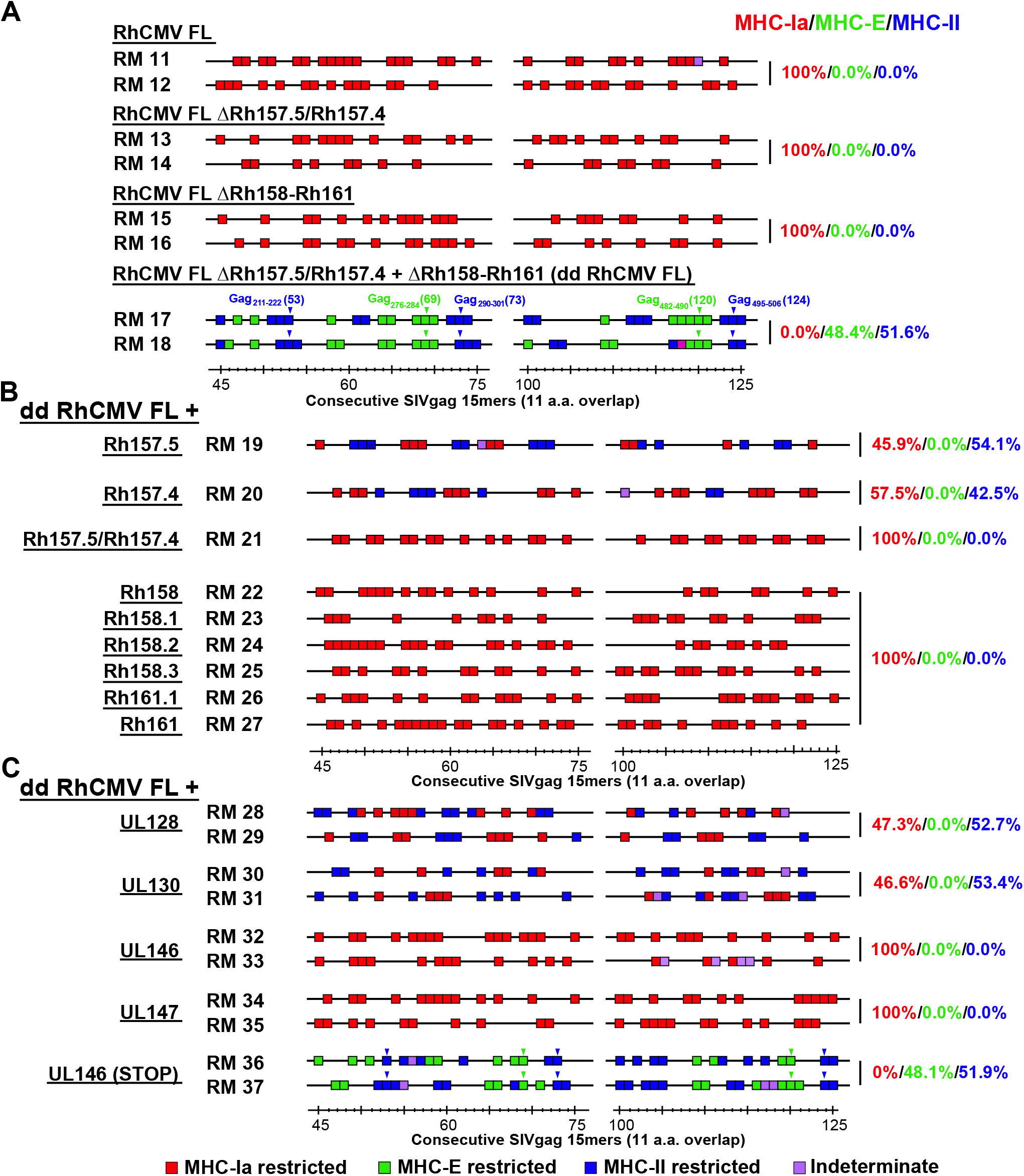
Identification of non-PRC regulators of RhCMV vector CD8^+^ T cell response programming. **(A-C)** MHC restriction analysis of SIV-specific CD8^+^ T cell responses elicited by the SIVgag-expressing FL RhCMV (wildtype configuration) vector and the designated modifications of this vector (see **fig. S1**). These analyses were identical to those in **Fig. 1** and are presented as described in **Fig. 1** legend. Overall epitope analysis results are presented in **table S1**.

To determine the effect of each of the eight genes missing from dd RhCMV FL (Rh157.5, Rh157.4; Rh158, Rh158.1, Rh158.2, Rh158.3, Rh161.1, Rh161) on CD8^+^ T cell response programming, we engineered their individual expression in the Rh161 locus of the dd RhCMV FL vector (**fig. S1**). We also evaluated adding back dual expression of the PRC components Rh157.5 and Rh157.4 to this vector. Adding back expression of Rh157.5 alone or Rh157.4 alone to dd RhCMV FL resulted in CD8^+^ T cell targeting of both MHC-Ia + MHC-II-restricted epitopes, whereas adding expression of both resulted in MHC-Ia-only epitope targeting (**Fig. 2B**), independently confirming the findings reported above using the 68-1.2, 68-1.2 ΔRh157.5, and 68-1.2 ΔRh157.4 vectors. In contrast, adding back expression of each of the six Rh158-161 genes individually to dd RhCMV FL resulted in reversion to the MHC-Ia-only epitope targeting pattern (**Fig. 2B**). Although only three of these genes are completely deleted in 68-1 RhCMV (**fig. S1**), qPCR analysis indicates that the remaining three transcripts are expressed at substantially lower levels in 68-1 compared to RhCMV FL, presumably as a consequence of the inversion event resulting in altered transcriptional regulation or ORF truncation (**fig. S5**). Taken together, these data indicate that wildtype RhCMV encodes highly redundant mechanisms to prevent elicitation of unconventionally restricted CD8^+^ T cell responses, particularly MHC-E-restricted CD8^+^ T cells. These include eight chemokine-related gene products (**fig. S4**), six of which (Rh158-161) that inhibit both MHC-E and MHC-II-restricted CD8^+^ T cell priming, and two of which (Rh157.5 and Rh157.4) that individually inhibit MHC-E- and MHC-II supertope CD8^+^ T cell priming and that when expressed together with UL131, gH, and gL create the PRC which additionally inhibits MHC-II non-supertope CD8^+^ T cell priming by tropism modulation.

Since our ultimate aim is to develop an HCMV-based vaccine for human use (*19*), we next asked whether the functions that are inhibitory to unconventional response generation are conserved in the HCMV orthologs of these gene products. As described for the RhCMV genes above, dd RhCMV FL vectors were engineered to individually express HCMV UL128, UL130, UL146, and UL147. Strikingly, these HCMV orthologs, though considerably sequence diverse from the RhCMV counterparts (**fig. S4**), showed the same pattern of inhibition of unconventional CD8^+^ T cell priming exhibited by the RhCMV gene products (**Fig. 2C**). Of note, stop mutations abrogated the inhibitory activity of HCMV UL146, consistent with a function of the UL146 protein being responsible for the observed inhibitory activity (**Fig. 2C**). This remarkable conservation of function strongly suggests that unconventional CD8^+^ T cell response inhibition is also a priority for HCMV infection in humans, and that deletion of at least these 4 genes (UL128, UL130, UL146, UL147) from HCMV would be required to elicit unconventionally restricted CD8^+^ T cells. These observations likely explain the lack of unconventionally restricted CD8^+^ T cell responses in humans vaccinated with a PRC negative, but UL130-, UL146- and UL147-expressing Towne/Toledo chimeric HCMV (*20*).

The ability of 68-1 RhCMV to elicit unconventionally restricted CD8^+^ T cell responses thus resulted from a fortuitous genetic event that precisely abrogated the activity of 8 genes in two non-contiguous, multi-gene regions. These data, however, do not indicate whether unconventional CD8^+^ T cell response priming, though present in the 68-1 RhCMV/SIV vectors with demonstrated efficacy (*6–8*), is required for, or associated with efficacy. To address this question, we initiated a vaccine-challenge study to compare the T cell immunogenicity and efficacy of RhCMV/SIV vaccines (vector sets comprised of 3 vectors of the same genetic configuration, individually expressing SIVgag, retanef, and 5’-pol) that elicit CD8^+^ T cell responses of the three identified CD8^+^ T cell targeting response types: 1) MHC-E + MHC-II, with supertopes (68-1), 2) MHC-Ia-only (68-1.2), and 3) MHC-Ia + MHC-II, no supertopes (68-1.2 ΔRh157.5) (**fig. S6**). To ascertain whether a vaccine regimen eliciting CD8^+^ T cell responses targeting a broad combination of all types of CD8^+^ T cell epitopes (MHC-Ia- and MHC-E- and MHC-II-restricted, including supertopes) was possibly more efficacious than individual vector sets, we also studied an additional 15 RM that were vaccinated with both the 68-1 and 68-1.2 vector sets.

Longitudinal analysis of SIV-specific CD4^+^ and CD8^+^ T cell responses from these RM cohorts showed that the 68-1.2, 68-1.2 ΔRh157.5, 68-1+68-1.2 vaccines manifested equivalent or greater SIV insert-specific T cell response magnitudes and similar response longevity as the 68-1 reference vaccine in both blood and bronchoalveolar lavage fluid (BAL, representing an effector site) (**Figs. 3A,B; fig. S7**). Epitope restriction analysis revealed the expected MHC restriction patterns for all four RhCMV/SIV vaccines, including supertope recognition only in the RM vaccinated with 68-1 vectors (**Figs. 3C; fig. S8**). Importantly, in all vaccine groups, the plateau phase SIVgag-specific CD4^+^ and CD8^+^ T cells manifested the characteristic EM-bias of RhCMV vector-elicited T cell responses, as well as broadly similar cytokine profiles, indicating that the genetic differences between these vectors did not impact the unique characteristics of RhCMV-induced T cell immunogenicity (**Figs. 3D,E**). We also compared the magnitude of SIVgag-specific CD4^+^ and CD8^+^ T cell responses in tissues of RMs vaccinated with 68-1 vs. 68-1.2 RhCMV/gag vectors studied at necropsy, revealing comparable responses across lymphoid and non-lymphoid tissues (**fig. S9**). Taken together, these data indicate that while these vaccines elicit SIV-specific CD8^+^ T cells with different epitope targeting, the magnitude, longevity, differentiation and tissue distribution of these CD8^+^ T cells, and of the SIV-specific CD4^+^ T cells, are broadly similar, allowing assessment of the contribution of unconventional response targeting to protective efficacy.

**Figure 3.**
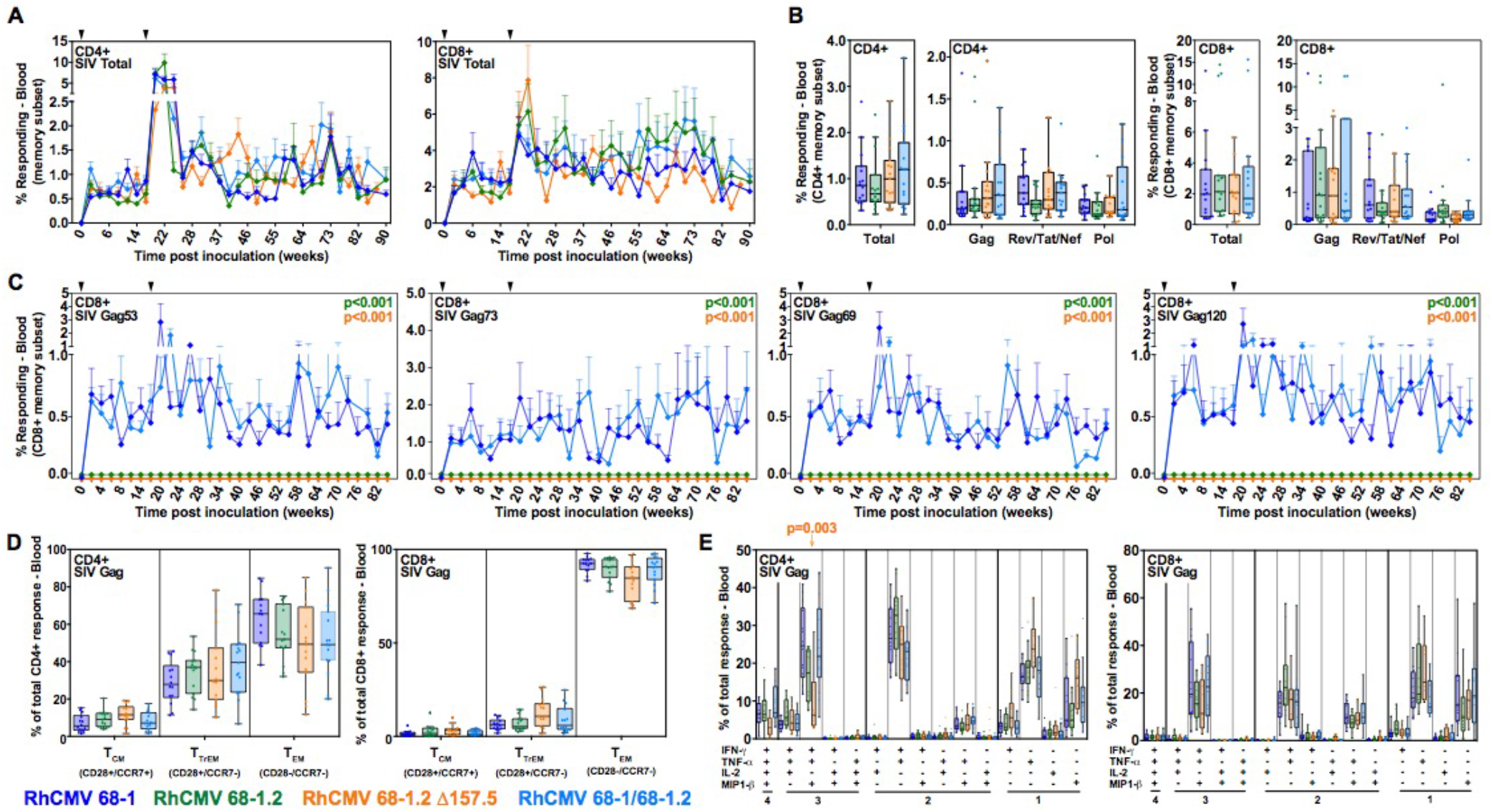
Immunogenicity of differentially programmed RhCMV vectors. **(A,B)** Longitudinal and plateau-phase analysis of the vaccine-elicited SIV Gag-, Rev/Tat/Nef-, and 5’-Pol-specific CD4^+^ and CD8^+^ T cell responses in peripheral blood of the RM vaccinated with 68-1, 68-1.2, and 68-1.2 ΔRh157.5 RhCMV vector sets and the combination of the 68-1 and 68-1.2 RhCMV vector sets (each set comprised of 3 vectors individually expressing SIV Gag, Rev/Tat/Nef, and 5’-Pol inserts; n = 15 RM per group). In **A**, the background-subtracted frequencies of cells producing TNF and/or IFN-γ by flow cytometric ICS assay to peptide mixes comprising each of the SIV inserts within the memory CD4^+^ or CD8^+^ T cell subsets were summed for overall responses with the figure showing the mean (+ SEM) of these overall responses at each time point (area-under-the-curve was used to quantitatively compare longitudinal response profiles). In **B**, boxplots compare the total and individual SIV insert-specific CD4^+^ and CD8 ^+^ T cell response frequencies between the vaccine groups during the vaccine phase plateau (each data point is the mean of response frequencies in all samples from weeks 61-90 post-first vaccination). (**C**) Longitudinal analysis of the vaccine elicited CD8^+^ T cell responses to SIV Gag supertopes in peripheral blood of each vaccine group by ICS assay. Gag_276-284_ (69) and Gag_482-490_ (120) are MHC-E-restricted supertopes; Gag_211-222_ (53) and Gag_290-301_ (73) are MHC-II-restricted supertopes. (**D**) Boxplots compare the memory differentiation of the vaccine-elicited CD4^+^ and CD8^+^ memory T cells in peripheral blood responding to overall SIV Gag 15mer peptide mix with TNF and/or IFN-γ production during the vaccine phase plateau (24-85 weeks post-first vaccination). Memory differentiation state was based on CD28 and CCR7 expression, delineating central memory (T_CM_), transitional effector-memory (T_TrEM_), and effector-memory (T_EM_), as designated. (**E**) Boxplots compare the frequency of vaccine-elicited CD4^+^ and CD8^+^ memory T cells in peripheral blood responding to the overall SIV Gag 15mer peptide mix with TNF, IFN-γ, IL-2, and MIP-1β production, alone and in all combinations, in the same samples as panel D. Wilcoxon p-values for comparison of all response parameters shown in panels A-E for the 68-1-only vaccine to all other vaccines are shown where significant (adjusted for multiple comparisons in panels B-E).

Starting ninety-one weeks after first vaccination, all vaccinated RM cohorts, and an unvaccinated RM cohort (n = 15), were subjected to repeated, limiting dose, intrarectal SIVmac239 challenge, with take of infection monitored by *de novo* induction of SIVvif-specific T cell responses (SIVvif is not present in the vaccine inserts and thus such responses are derived from the SIV infection) and plasma viral load (pvl), as previously described (*6–8*). In this well-characterized challenge model, unvaccinated controls and non-protected RM manifest typical peak and plateau-phase SIVmac239 pvl in concert with development of SIVvif-specific T cells, whereas protected RM show development of SIVvif-specific responses in the absence of plasma viremia or with non-sustained pvl blips, and manifest cell-associated SIV DNA and/or RNA in tissues consistent with replication arrest (*6–8*). As expected, all unvaccinated RMs showed typical progressive infection, whereas 8 of 15 (53%) of the 68-1 vector vaccinated RMs showed protection (**Figs. 4A,B**). However, none of the RMs vaccinated with the 68-1.2 vectors or 68-1.2 ΔRh157.5 vectors manifested protection, but notably, protection was still observed in 6 of the 15 (40%) RMs vaccinated with both 68-1 and 68-1.2 vectors (p = NS relative to 68-1) (**Figs. 4C-E**). Protected RM in both the 68-1 and 68-1 + 68-1.2 RhCMV/SIV vector vaccinated groups manifested cell-associated SIV DNA and/or RNA in bone marrow (BM) and lymph node (LN), confirming take of infection, and given the absence of measurable viremia, replication arrest (**Fig. 4F**). Thus, although the RhCMV vectors programmed for the alternative CD8^+^ T cell targeting response types (MHC-Ia-+ MHC-II-restricted) elicited CD8^+^ T cell responses with similar magnitude and qualitative characteristics as 68-1 vectors, these responses were unable to provide protection against SIV challenge.

**Figure 4.**
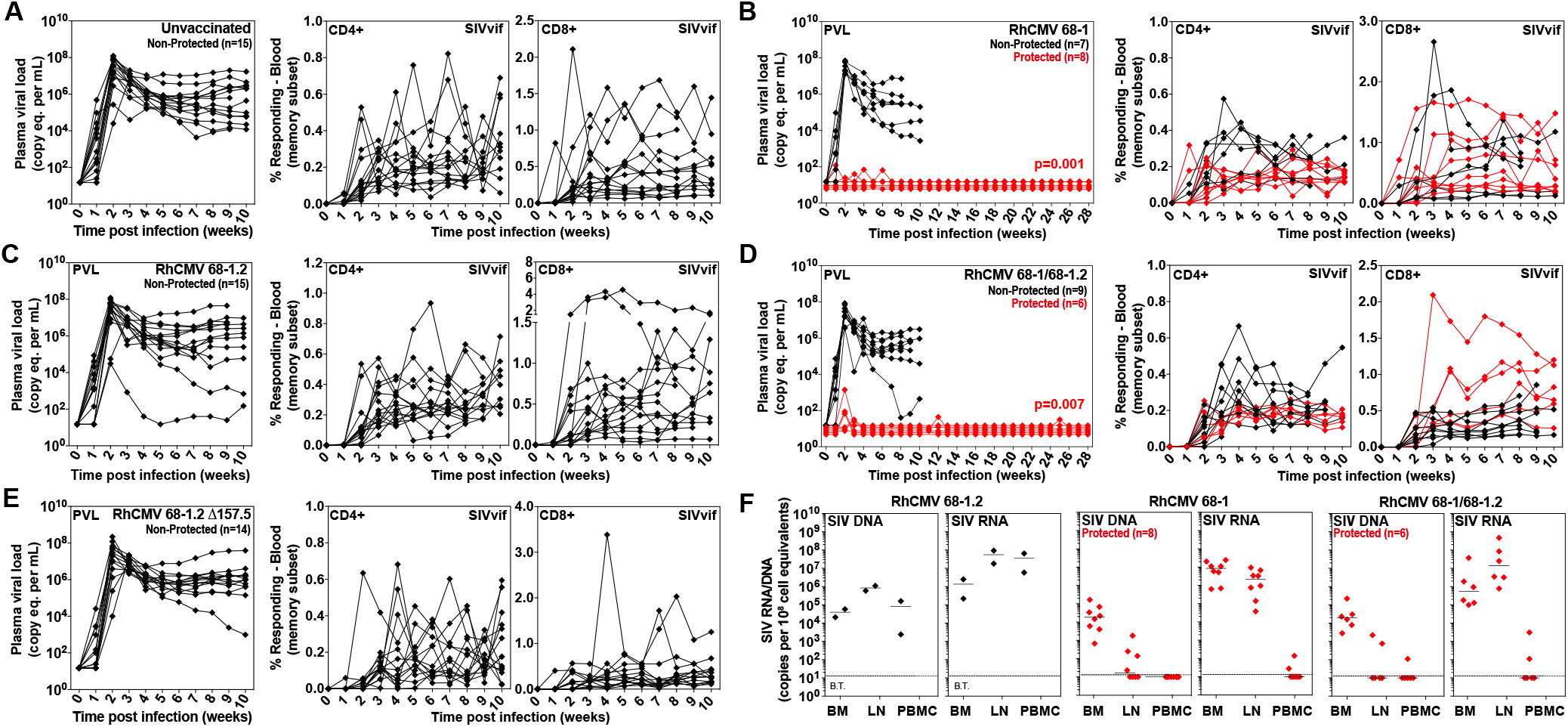
Efficacy of differentially programmed RhCMV vectors. **(A-E)** Assessment of the outcome of SIV infection after repeated, limiting dose SIV_mac239_ challenge of the designated vaccine groups by longitudinal analysis of plasma viral load (left panels) and *de novo* development of SIVvif-specific CD4^+^ (middle panels) and CD8^+^ (right panels) T cell responses. RM were challenged until the onset of any above-threshold SIVvif-specific T cell response, with the SIV dose administered 2 or 3 weeks prior to this response detection considered the infecting challenge (week 0). RM with sustained viremia were considered non-protected (black); RM with no or transient viremia, but clear SIVvif-specific T cell responses, were considered protected (red) (*6–8*). Binomial exact p-values are shown where the proportion of protected RM in a vaccine group differs significantly from the unvaccinated group. (**F**) Bone marrow (BM), peripheral lymph node (LN) and peripheral blood mononuclear cell (PBMC) samples from all vaccine-protected RM (red) and representative unprotected, control RM (black), collected from between day 28 and 56 post-SIV infection, were analyzed by nested, quantitative PCR/RT-PCR for cell-associated SIV DNA and RNA. The dotted line indicates the threshold of detection (B.T. = below threshold) with data points below this line reflecting no positive reactions across all replicates.

In a companion manuscript (*21*), we demonstrate that 68-1 vectors deleted for Rh67 (RhCMV ortholog of HCMV UL40), the gene product which is responsible for MHC-E upregulation in vector-infected cells, specifically fail to prime MHC-E-restricted CD8^+^ T cells, leaving CD8^+^ T cell responses which are only MHC-II-restricted, including MHC-II-restricted supertopes. Taken together, these data provide unequivocally evidence that RhCMV has evolved the ability to control the nature of its own recognition by CD8^+^ T cells – to our knowledge, the only virus with this capability, and thus the only virus-based vaccine vector that is genetically programmable with respect to CD8^+^ T cell epitope targeting. Importantly, the exclusively MHC-II-restricted SIV-specific CD8+ T cells responses elicited by the Rh67-deleted 68-1 RhCMV/SIV vectors, though also of similar (or higher) magnitude, longevity and effector memory differentiation as 68-1 RhCMV/SIV vector elicited responses, were, like the MHC-Ia- and MHC-Ia + MHC-II-restricted CD8^+^ T cells elicited by 68-1.2 and 68-1.2 ΔRh157.5 vectors in this study, unable to provide protection against SIV challenge (*21*). CD4^+^ T cell responses are comparable across these differentially programmed RhCMV/SIV vaccines and antibody responses are largely absent in 68-1 RhCMV/SIV vaccinated RM (*6–8, 22*), which makes the presence of MHC-E-restricted CD8^+^ T cell responses the only defined difference in adaptive (SIV-specific) immunity between efficacious and non-efficacious RhCMV/SIV vectors. Since SIV inserts (and thus SIV-specific immunity) are demonstrably required for efficacy (*7*), these data strongly suggest that the MHC-E-restricted CD8^+^ T cell response is the crucial adaptive immune component RhCMV/SIV vector immunogenicity that mediates “control and clear” protection. Although the specific immune functions mediating this efficacy *in vivo* remain to be elucidated (in particular, delineation of why MHC-E-restricted CD8^+^ T cell response are uniquely efficacious), these data provide the first documentation of a vaccine for CD8^+^ T cell immunity that includes programmability for unconventionally restricted CD8^+^ T cell responses with unique *in vivo* efficacy. The finding that the HCMV orthologs of the RhCMV genes that mediate CD8^+^ T cell response programming, including UL128/UL130, UL146/UL147 and UL40 (*21*), recapitulate the response programming function of their RhCMV counterparts when used to replace these counterparts in RhCMV strongly suggests the conservation of CD8^+^ T cell response programming in HCMV and supports the feasibility of translating this unique immunobiology to humans to create an effective HIV/AIDS vaccine. The flexible programmability of CMV vectored vaccines may also be exploitable in development of vaccines to other pathogens or to malignancies.

## ACKNOWLEDGMENTS

We thank T. Whitmer, A. Bhusari, L. Bishop, C. Kreklywich, A. Legasse, M. Fischer, C. Shriver-Munsch, T. Swanson, A. Sylwester, S. Hagen, E. McDonald, K. Randall, A. Selseth, and K. Rothstein for technical or administrative assistance; B. Keele for providing SIVmac239 challenge virus, and both J. Womack and A. Townsend for figure preparation.

## AUTHOR CONTRIBUTIONS

DM, DS, and KF designed, constructed and validated the modified 68-1, 68-1.2 and FL RhCMV vectors used in this study (assisted by LSU, MRM and CRP), except the miR-142-p3 tropism-restricted 68-1.2 RhCMV vector which was conceived, constructed and validated by MHH, FG and JAN, assisted by RET. SGH planned and performed animal experiments and immunologic assays, assisted by CMH, JCF, DM, RMG, ABV, JBS and JMG. MKA managed the animal care and procedures. JDL planned and supervised SIV quantification by PCR/RT-PCR assisted by KO, RS, BB, WJB and MH. AWL and. JS and PTE conceived and performed statistical analyses. LJP conceived the RhCMV vector strategy, supervised all experiments, analyzed and interpreted data, and wrote the paper, assisted by SGH, DM, JBS, JDL, PTE, and KF.

## DATA AND MATERIALS AVAILABILITY

All data associated with this study are present in the paper or Supplementary Materials. The computer code used to perform statistical analysis is available upon request. RhCMV/SIV vectors can be obtained through an MTA.

