## Supplemental Material for "Cytomegaloviral determinants of CD8^+^ T cell programming and RhCMV/SIV vaccine efficacy"

### SUPPLEMENTARY MATERIALS

#### Materials and Methods

**Figure S1.** Genetic configuration of RhCMV vectors.

**Figure S2.** Functional PRC analysis of 68-1.2-based RhCMV vectors.

**Figure S3.** Myeloid cell tropism-restriction of a 68-1.2 RhCMV with microRNA (miR)-142.

**Figure S4.** Protein sequence comparison of RhCMV/HCMV ortholog RhCMV157.5/UL128, Rh157.4/UL130 and Rh158-161/UL146-147

**Figure S5.** Transcriptional analysis of Rh158-Rh161 region genes in RhCMV FL vs. 68-1/68-1.2.

**Figure S6.** Protocol for the comparison of the immunogenicity and efficacy of 68-1, 68-1.2, and ΔRh157.5 68-1.2 RhCMV/SIV vector sets, and the combination of 68-1 and 68-1.2 vectors.

**Figure S7.** Analysis of SIV-specific T cell responses in BAL in RM cohorts destined for SIV challenge.

**Figure S8.** Epitope analysis of RhCMV vaccine-elicited, SIV-specific CD8+ T cell responses in RM cohorts destined for SIV challenge.

**Figure S9**. Analysis of systemic SIVgag-specific CD4+ and CD8+ T cell response magnitude in 68-1 vs. 68-1.2 RhCMV/SIVgag vector vaccinated RMs at necropsy.

**Table S1.** Epitope analysis summary of all study RMs.

### MATERIALS AND METHODS

**Study Design.** This study had two major objectives. First, we sought to determine the molecular genetic basis of unconventional CD8⁺ T cell response programming by strain 68-1 RhCMV vectors, knowing that naturally occurring RhCMVs elicit conventional MHC-Ia-restricted CD8⁺ T cells, whereas the genetically distinct 68-1 RhCMV elicits a mixture of MHC-II- and MHC-E-restricted CD8⁺ T cells (*9, 10*). We had previously identified the complex inversion/deletion in the genomic region of RhCMV orthologous to the ULb' region of HCMV as the likely basis of this immune programming (**Fig. 1A**), but the specific contribution of the affected genes was unclear, as well as the spectrum of immunologic phenotypes (response types) that might arise from different configurations of this gene region. Once the genetic determinants of unconventional CD8⁺ T cell response priming were identified and the spectrum of different response types established, our second major objective was to both compare the overall immunogenicity of all distinct RhCMV response types and to determine whether the RhCMV vectors eliciting these alternative response types were able to mediate the unique replication arrest form of protective efficacy previously described for 68-1 RhCMV/SIV vectors (*6-8*). We sought to determine the basis of 68-1 RhCMV/SIV vector efficacy, in particular the dependence of this efficacy on unconventional CD8⁺ T cell epitope targeting.

Our approach to Objective #1 was to 1) construct SIVgag-expressing RhCMV vectors with defined deletions, additions, or functional inactivations of specific open reading frames (ORFs) in the ULb' ortholog region (**Figs. 1A; fig. S1**), 2) strategically vaccinate monkeys with these variants, 3) determine the effect of each ORF modification on CD8⁺ T cell recognition of SIV epitopes (with 34 to >500 epitopes analyzed), and 4) confirm gene contributions and response types with multiple distinct, but functionally overlapping, vectors. For example, the distinct role of the PRC in response programming vs. PRC-independent effects of Rh157.5/UL128 and Rh157.4/UL130 gene products, a difference resulting in the unique mixed MHC-Ia + MHC-II (no supertope) response type, was documented in 15 RM vaccinated with the 68-1.2 ΔRh157.5 vector, 4 RM vaccinated with the 68-1.2 ΔRh157.4 vector, 2 RM vaccinated with 68-1.2 with the CC1+CC2 mutant-Rh157.5, 2 RM vaccinated with dd FL RhCMV with either Rh157.5 or Rh157.4 added back, 2 RM vaccinated with dd FL RhCMV with either HCMV UL128 or UL130 added back, and 3 RM vaccinated with the miR-142-39 tropism-restricted 68-1.2 RhCMV, which recapitulates a major tropism effect of PRC loss. All told 28 RM encompassing separate analysis of 974 peptide-specific responses, encompassing an estimated 666 distinct epitopes (**table S1**).

For Objective #2, we selected 3 vector backbones and 1 vector backbone combination for comparison, encompassing the 3 response types identified in our genetic modification studies (MHC-Ia, MHC-Ia + MHC-II, MHC-E + MHC-II with supertopes) and a combination phenotype that elicited all these responses. Based on previous experience with 68-1 RhCMV/SIV vectors (*6-8*), we randomly assigned male RM to one of the vaccine groups or to serve as unvaccinated controls (n=15 per group; n=75 total). This group size was anticipated to allow us to thoroughly compare overall immunogenicity of each vector backbone or combination and to detect per-vaccine-group protection levels of 14% at 90% power without multiplicity adjustment. All results from these experiments are included in the presented associated data (no data were excluded as outliers). Plasma and cell-associated viral load assays were performed by blinded analysis; however, due to logistical constraints, other staff were not blinded to treatment assignments. Primary data are reported in data files 1-3.

**Rhesus macaques.** These experiments used a total of 124 purpose-bred male and female RM (*M. mulatta*) of Indian genetic background, including 1) 43 RM for immunogenicity analysis of gene-modified 68-1, 68-1.2 and FL RhCMV/gag vectors, 2) 75 RM for immunogenicity and efficacy analysis of RhCMV vector backbones representing distinct response types (5 groups of n=15, 4 groups vaccinated; 1 group unvaccinated), and 3) 6 RM for RhCMV 68-1.2 vs. 68-1 vector immunogenicity analysis at necropsy. Of note, the 68-1 vaccine group from this study serves as a positive control group for the companion manuscript (*21*). RhCMV vectors were routinely dosed at $10^6$-$10^7$ infectious units for immunogenicity analysis and $5x10^6$ infectious units per vector for efficacy analysis, all via subcutaneous administration. At assignment, all study RM were free of cercopithicine herpesvirus 1, D-type simian retrovirus, simian T-lymphotrophic virus type 1, and *Mycobacterium tuberculosis*, but all except 1 (RM38 in **fig. S9**) were naturally RhCMV-infected. All study RM were housed at the Oregon National Primate Research Center (ONPRC) in Animal Biosafety level (ABSL)-2 (vaccine phase) and ABSL-2+ rooms (challenge phase) rooms with autonomously controlled temperature, humidity, and lighting. Study RM were both single and pair cage housed. Animals were only paired with one another during the vaccine phase, if they were from the same vaccination group. All RM were single cage-housed during the challenge phase due to the infectious nature of the study. Regardless of their pairing, all animals had visual, auditory and olfactory contact with other animals. Single cage-housed RM received an enhanced enrichment plan that was designed and overseen by NHP behavior specialists. RM were fed commercially prepared primate chow twice daily and received supplemental fresh fruit or vegetables daily. Fresh, potable water was provided via automatic water systems. Physical exams including body weight and complete blood counts were performed at all protocol time points. RM were sedated with ketamine HCl or Telazol for procedures, including intradermal and subcutaneous vaccine administration, venipuncture, bronchoalveolar lavage, BM and LN biopsy, and SIV challenge. At humane or scheduled endpoints, RM were euthanized with sodium pentobarbital overdose (>50 mg/kg) and exsanguinated via the distal aorta, and tissue collection at necropsy was performed by a certified veterinary pathologist. RM care and all experimental protocols and procedures were approved by the ONPRC Institutional Animal Care and Use Committee (IACUC). The ONPRC is a Category I facility. The Laboratory Animal Care and Use Program at the ONPRC is fully accredited by the American Association for Accreditation of Laboratory Animal Care (AAALAC) and has an approved Assurance (#A3304-01) for the care and use of animals on file with the NIH Office for Protection from Research Risks. The IACUC adheres to national guidelines established in the Animal Welfare Act (7 U.S.C. Sections 2131–2159) and the Guide for the Care and Use of Laboratory Animals (8th Edition) as mandated by the U.S. Public Health Service Policy.

**Generation of recombinant RhCMV vectors.** SIV insert (SIVgag, retanef, 5'pol)-expressing vaccine vectors based on RhCMV 68-1 (GenBank #MT157325) have been described previously (*6, 23*). All other RhCMV constructs used in this study (**fig. S1**) were based on either the partially-repaired RhCMV 68-1.2 strain (#MT157326) or our fully reconstructed FL RhCMV clone (#MT157327) (*13, 18*). Following the original 68-1 vector design, SIV inserts were introduced into the Rh211 open reading frame (ORF) in RhCMV 68-1.2 and transgene expression was driven by an EFIα promoter. For FL-RhCMV vectors, this design was altered and SIV transgenes were inserted into the Rh13.1 ORF using the endogenous promoter and regulatory elements to drive expression. This configuration resulted in increased genome stability as Rh13.1 is selected against in low passage isolates during tissue culture (*18*). To introduce deletions into these BACs using recombineering we either chose a classical lambda (λ) Red recombination system in combination with flippase recognition target (FRT) sites (*24*), *en passant* recombination (*25*) or galactokinase- (galK-) mediated BAC recombination (*26*). Red recombination is performed by amplifying an aminoglycoside 3-phosphotransferase gene conferring kanamycin resistance (KanR) flanked by FRT sites with primers carrying a 50bp homology arm to the upstream and downstream region of the targeted genomic locus. This PCR product is inserted into the BAC through homologous recombination by heat shock induced expression of the lambda prophage encoded red recombination genes. After XmaI restriction digest and Sanger sequencing across the altered sequence, the selection marker is removed through the arabinose inducible expression of a flip recombinase in the EL250 or SW105 strains of *E. coli*. Final constructs were analyzed by restriction digest and Sanger sequencing of the altered genomic regions. Unintended off-target alterations in the genome sequence were ruled out by next generation sequencing (NGS) of the complete genome using an Illumina MiSeq or iSeq sequencing platform.

The *en passant* recombination technique utilizes a strategy which includes the introduction of 50bp DNA repeats during primer design flanking the KanR selection marker. An I-SceI homing enzyme target sequence is introduced between the selection marker and the 5'homology arm, which can be used to selectively introduce DNA double strand breaks after arabinose inducible expression of the homing enzyme in *E. coli* strain GS1783. These breaks can be repaired via recombination of the repeated DNA sequences flanking the selection marker by inducing the expression of the lambda phage derived Red recombination genes through heat shock, resulting in complete removal of the KanR resistance cassette without retaining any DNA sequences introduced during recombineering (scarless removal). Hence, the latter technique was favored for the introduction of point mutations or the alteration of neighboring amino acids so as to not disturb the overall coding region. We also used this technique to introduce HCMV TR3 (#MN075802) or RhCMV FL derived ORFs into the Rh161 locus in dd RhCMV FL using the endogenous regulatory elements to drive gene expression. As described for the lambda Red recombination system, all constructs were analyzed by XmaI restriction digest, Sanger sequencing of the altered genome region as well as full genome analysis by NGS before virus reconstitution.

The RhCMV 68-1.2-miR-142-3p and 68-1.2-scrambled miR control vectors expressing SIVgag were constructed using galactokinase- (galK-) mediated BAC recombination. The SW105 *E. coli* strain carrying the 68-1.2 SIVgag BAC was used to introduce an expression cassette encoding the galactokinase as well as the KanR gene flanked by 80bp homology arms to the targeted regions in the Rh156 or Rh108 3' untranslated regions. Correctly recombined clones were identified based on the production of bright pink colonies on MacConkey agar containing kanamycin and Sanger sequencing of the inserted region. The galK-KanR cassette was replaced by the insert of interest by homologous recombination and correctly recombined clones were identified through negative selection on 2-deoxy-galactose-containing plates. These constructs were also analyzed by restriction digest, Sanger sequencing of the inserted region as well as full genome analysis by NGS before virus reconstitution.

**RhCMV vector recovery and characterization.** All vectors derived from BACs were reconstituted in primary embryonal rhesus fibroblasts (RFs) via electroporation (250V, 508 950μF) of purified BAC-DNA. The cells were maintained in DMEM complete, Dulbecco's modified Eagle's medium (DMEM) with 10% fetal bovine serum and antibiotics (1× Pen/Strep; Gibco), and grown at 37°C in humidified air with 5% $CO_2$. Depending on the reconstituted construct, viral plaques became visible within 3-5 days and full cytopathic effect (CPE) was observed within 1-2 weeks after electroporation. At this point, cells and supernatants were harvested and stored at -80°C until final use. Viral stocks were produced on RFs by infecting eight confluent T-175 tissue culture flasks with an approximate MOI of 0.05 – 0.1. After full CPE was reached within 4-8 days cells and supernatants were harvested and frozen once overnight at -80°C to release intracellular virus from infected cells. Subsequently the supernatant was clarified by centrifugation in two steps, first at 2,000 x g for 10 minutes at 4°C and secondly at 7,500 x g for 15 minutes, after which the virus was purified through a sorbitol cushion (20% D-sorbitol, 50 mM Tris [pH 7.4], 1 mM $MgCl_2$) by centrifugation at 64,000 x g for 1 hour at 4°C in a Beckman SW28 rotor. The pelleted virus was resuspended in complete DMEM, aliquoted and stored at -80°C until use. Virus titers of all RhCMV stocks were determined by fifty-percent tissue culture infective dose ($TCID_{50}$) assay on RFs and expression of the utilized SIV immunological marker was confirmed by immunoblot.

To determine which RhCMV constructs contained a functional PRC, we compared entry into rhesus retinal pigment epithelial cells (RPE) to entry into primary rhesus fibroblast (RF). RPEs were a kind gift from Dr. Thomas Shenk (Princeton University, USA) and were propagated in a 1:1 mixture of DMEM and Ham's F12 nutrient mixture with 5% FBS, 1 mM sodium pyruvate, and nonessential amino acids. Both RFs and RPEs were infected with a MOI determined on RF resulting in infection levels of 20-30 percent after 48 hours of infection (MOI of 0.3 for 1RFs and MOI of 10 for RPEs) using the PRC intact RhCMV strain 68-1.2 as a positive control. Infection levels were determined by flow cytometry using a RhCMV specific antibody (*27*) and infection levels were expressed by setting infection in RFs to 100 per cent and expressing infection levels in RPEs in relation to infection levels in RFs. RhCMV strains 68-1 and 68-1.2 were included in all assays as negative and positive controls, respectively, and all experiments were performed with triplicate repeats.

To analyze the growth restriction of the RhCMV-miR-142-3p vector, multi-step growth curve assays were performed on primary rhesus fibroblasts transiently transfected with miR-142-3p or a negative control mimic. 12-well plates seeded with RFs were transfected with 20pmol miRNA mimic/well (Dharmacon) using RNAiMax (Invitrogen) according to the manufacturer's instructions. 24 hours later the cells were infected with RhCMV-miR-142-3p or the scrambled control virus at a multiplicity of infection of 0.01. Supernatants were collected at the indicated timepoints and titered using a standard plaque assay on RFs. To analyze growth restriction in primary rhesus macrophages, rhesus PBMC were isolated from whole blood using Ficoll and monocytes were purified using non-human primate $CD14^+$ microbeads (Miltenyi). The isolated monocytes were cultured in 50% complete RPMI supplemented with 10% FBS, 18ng/mL human GM-CSF (R&D Systems) and 20ng/mL human M-CSF (R&D Systems) and 50% conditioned media from the cell line KPB-m15 (2) for 7 days on Primaria plates (ThermoFisher Scientific). One half media exchange occurred on days 2, 4 and 6. After this time cells had adhered to the Primaria plates and differentiated to a typical macrophage morphology. Macrophages were infected with RhCMV 68-1.2 miR-142-3p or scrambled control vector at MOI = 5 and cell lysates was harvested at the indicated times post-infection using Trizol (Invitrogen). DNA was isolated and used to perform quantitative PCR (qPCR) for RhCMV genomes using custom primers and probe sets (ThermoFisher Scientific) for RhCMV Rh156: RhCMV Rh156 F primer: GGGCATCCTCAGGATCACAG; RhCMV Rh156 R primer: CGACACCAAGAGGGTATGGG; RhCMV Rh156 probe: 6FAM-ACTCCGAAGACCACAAGGACCCACG-BHQ1. Standard curves were prepared using RhCMV BAC DNA.

**Expression analysis of RhCMV encoded inhibitors of unconventionally restricted $CD8^+$ T-cell priming.** Primary rhesus fibroblasts were seeded out in tissue culture plates and infected the next day with either RhCMV FL, 68-1, or 68-1.2 at a MOI of 5. Mock infected wells were included as negative control samples. The cells were harvested at 0, 4, 8, 12, 24, 36, and 48 hours post infection (hpi) and total RNA was isolated using the Quick RNA Microprep kit (Zymo Research) according to the manufacturer's instructions. 1 µg of total RNA per sample was transcribed into cDNA using the Maxima Reverse Transcriptase (ThermoFischer Scientific). To examine the transcript levels and kinetics of selected RhCMV ORFs, a q-PCR assay was applied to the cDNA using TaqMan Fast Advanced Master Mix (Applied Biosystem). qPCR reactions were performed using QuantStudio 7 Flex Real-Time PCR Systems (Applied Biosystems) and data were collected using the QuantStudio Real-Time PCR Software v1.3. All determined transcript copy numbers were normalized to the housekeeping gene (GAPDH) and results for each gene and time point are expressed as relative mRNA copy numbers. As controls for our kinetic class analysis, we included the known immediate early (IE) gene Rh156 (UL123, IE1), the characterized early (E) gene Rh189 (US11) as well as the described true late (L) gene Rh137 (UL99, pp28). Each forward and reverse primer set was used initially to generate a PCR fragment specific for each gene which was cloned into the pGEM-T Easy Vector (Promega) according to the manufacturer's instructions. These plasmids were used for the generation of primer/probe set specific standard curves for each indicated gene which were required to calculate viral genome copy numbers. The primer/probe sets used in this study are as follows:

| | | |
|---|---|---|
| **Rh156 (IE1)** | Forward | 5' AGTATGCCAAGCCTCATATTAAGGA 3' |
| | Reverse | 5' GCATATGGTGCTTGCTCTTAGAAG 3' |

| | | |
|---|---|---|
| | Probe | 6FAMAAGGTGCTGGACCCMGBNFQ |
| **Rh189 (E)** | Forward | 5' GGAGCGCCCGGTAAGG 3' |
| | Reverse | 5' CGATGGAGTTTTATGCTTTGCA 3' |
| | Probe | 6FAMTCCAGTGCATCTGAATCMGBNFQ |
| **Rh137 (L)** | Forward | 5' GGCGCAACATACTACCCAGAA 3' |
| | Reverse | 5' GTAGCCATCCCCATCTTCCA 3' |
| | Probe | 6FAMCACAACTAACTCTGGCCTTMGBNFQ |
| **Rh157.5** | Forward | 5' CTCAACTACCTCAGGTCAATGTGACT 3' |
| | Reverse | 5' CAGGCGTTGGTGGCATAGTA 3' |
| | Probe | 6FAMTGCTACCTCTGCTTCATMGBNFQ |
| **Rh157.4** | Forward | 5' GCGAACGGCGAGATCAAC 3' |
| | Reverse | 5' TGGCTTCGGTGCCACTTG 3' |
| | Probe | 6FAMAACGCCAAGGTCAGCTMGBNFQ |
| **Rh157.6** | Forward | 5' AAAACACAAAACAACCCCATCTTAC 3' |
| | Reverse | 5' AATGTGATGAGTTTCTTTGCTGAAA 3' |
| | Probe | 6FAMCGTAGGATTGGATTAATTMGBNFQ |
| **Rh158** | Forward | 5' TCTACAACCACTACCCCCTAACG 3' |
| | Reverse | 5' TTTGGGAGCACAGCAATCAC 3' |
| | Probe | 6FAMAAATTGCACCAAACAAGAMGBNFQ |
| **Rh158.1** | Forward | 5' CGTAACCCGTCGGTGTTTG 3' |
| | Reverse | 5' GCTGCGGCTTCAAACTTATGTG 3' |
| | Probe | 6FAMATTGGATAAGCAAACATGMGBNFQ |
| **Rh158.2** | Forward | 5' ATAATCTCGGTTTTGTGACATTCACT 3' |
| | Reverse | 5' GCGTACCATGAAAGGCGTTAA 3' |
| | Probe | 6FAMTTTCACTATCTGAATATCTGMGBNFQ |
| **Rh158.3** | Forward | 5' CCCAAAACGATGGGTACGTT 3' |
| | Reverse | 5' GCCGATGATGAAGAAAATGTTG 3' |
| | Probe | 6FAMCTCCGGGAATTTTMGBNFQ |
| **Rh161.1** | Forward | 5' TCGGTTACTTCGCACCTAGGA 3' |
| | Reverse | 5' AGAATTCCACACAGGACACTATGC 3' |
| | Probe | 6FAMCCGCATATTCAATTGATMGBNFQ |
| **Rh161** | Forward | 5' AAAATCAGACTGTCTCTGGCTTCA 3' |
| | Reverse | 5' GGAAATGTGCTATTGCCTCGTTA 3' |
| | Probe | 6FAMAGTACGGCTCACCCTGTMGBNFQ |
| **GAPDH** | Forward | 5' TTCAACAGCGACACCCACTCT 3' |
| | Reverse | 5' GTGGTCGTTGAGGGCAATG 3' |
| | Probe | 6FAMCCACCTTCGACGCTGGMGBNFQ |

**Immunologic assays.** SIV-specific $CD4^+$ and $CD8^+$ T cell responses were measured in peripheral blood mononuclear cells (PBMC) or tissue-derived mononuclear cells by flow cytometric intracellular cytokine analysis, as previously described (*6-8*). Briefly, individual or whole protein mixes of sequential 15-mer peptides (11 amino acid overlap) spanning the $SIV_{mac239}$ Gag, 5'-Pol, Nef, Rev, Tat, and Vif proteins or individual $SIV_{mac239}$ Gag supertope peptides [$Gag_{211-222}$ (53), $Gag_{276-284}$ (69), $Gag_{290-301}$ (73), $Gag_{482-490}$ (120)] were used as antigens in conjunction with co-stimulatory anti-CD28 (CD28.2, Purified 500 ng/test: eBioscience, Custom Bulk 7014-0289-M050) and anti-CD49d mAb (9F10, Purified 500 ng/test: eBioscience, Custom Bulk 7014-0499-M050). Mononuclear cells were incubated at 37°C with individual peptides or peptide mixes and antibodies for 1 hr, followed by an additional 8 hr incubation in the presence of Brefeldin A (5 µg $ml^{-1}$; Sigma-Aldrich). Stimulation in the absence of peptides served as background control. After incubation, stimulated cells were stored at 4°C until staining with combinations of fluorochrome-conjugated monoclonal antibodies including: anti-CD3 (SP34-2: Alexa700; BD Biosciences, Custom Bulk 624040 and Pacific Blue; BD Biosciences, Custom Bulk 624034), anti-CD4 (L200: AmCyan; BD Biosciences, Custom Bulk 658025, BV510; BD Biosciences,

Custom Bulk 624340 and BUV395; BD Biosciences, Custom Bulk 624165), anti-CD8α (SK1: PerCP-eFluor710; Life Tech, Custom Bulk CUST04424), anti-TNF-α (MAB11: FITC; BD Biosciences, Custom Bulk 624046 and PE; BD Biosciences, Custom Bulk 624049), anti-IFN-γ (B27: APC; BD Biosciences, Custom Bulk 624078) and anti-CD69 (FN50: PE; eBioscience, Custom Bulk CUST01282 and PE-TexasRed; BD Biosciences, Custom Bulk 624005) and for polycytokine analyses, anti-IL-2 (MQ1-17H12; PE Cy-7; Biolegend), and anti-MIP-1β (D21-1351, BV421; BD Biosciences). For analysis of memory differentiation (central- vs transitional- vs effector-memory) of SIV Gag-specific $CD4^+$ and $CD8^+$ T cells, PBMC were stimulated as described above, except that the CD28 co-stimulatory mAb was used as a fluorochrome conjugate to allow CD28 expression levels to be later assessed by flow cytometry, and in these experiments, cells were surface-stained after incubation for lineage markers CD3, CD4, CD8, CD95 and CCR7 (see below for mAb clones) prior to fixation/permeabilization and then intracellular staining for response markers (CD69, IFN-γ, TNF-α; note that Brefeldin A treatment preserves the pre-stimulation cell-surface expression phenotype of phenotypic markers examined in this study).

Stained samples were analyzed on an LSR-II or FACSymphony A5 flow cytometer (BD Biosciences). Data analysis was performed using FlowJo software (Tree Star). In all analyses, gating on the lymphocyte population was followed by the separation of the $CD3^+$ T cell subset and progressive gating on $CD4^+$ and $CD8^+$ T cell subsets. Antigen-responding cells in both $CD4^+$ and $CD8^+$ T cell populations were determined by their intracellular expression of CD69 and either or both of the cytokines IFN-γ and TNF-α (or in polycytokine analyses, expression of CD69 and any combination of the cytokines: IFN-γ, TNF-α, IL-2, MIP-1β). Assay limit of detection was determined as previously described (*22*), with 0.05% after background subtraction being the minimum threshold used in this study. After background subtraction, the raw response frequencies above the assay limit of detection were "memory-corrected" (e.g., % responding out of the memory population), as previously described (*6-8, 22*), using combinations of the following fluorochrome-conjugated mAbs to define the memory vs naïve subsets CD3 (SP34-2: Alexa700 and PerCP-Cy5.5; BD Biosciences Custom Bulk 624060), CD4 (L200: AmCyan), CD8α (SK-1: PerCP-eFluor710, RPA-T8: APC; BioLegend), TNF-α (MAB11; FITC), IFN-γ (B27; APC), CD69 (FN50; PE), CD28 (CD28.2; PE/Dazzle 594, BioLegend), CD95 (DX2; PE, BioLegend), CCR7 (15053; Biotin, R&D Systems), streptavidin (Pacific Blue, Life Tech and BV605; BD Biosciences, Custom Bulk 624342) and Ki67 (B56; FITC, BD Biosciences, Custom Bulk 624046). For memory phenotype analysis of SIV Gag-specific T cells, all $CD4^+$ or $CD8^+$ T cells expressing CD69 plus IFN-γ and/or TNF-α were first Boolean OR gated, and then this overall Ag-responding population was subdivided into the memory subsets of interest on the basis of surface phenotype (CCR7 vs CD28). Similarly, for polycytokine analysis of SIV Gag-specific T cells, all $CD4^+$ or $CD8^+$ T cells expressing CD69 plus cytokines were Boolean OR gated and polyfunctionality was delineated with any combination of the four cytokines tested (IFN-γ, TNF-α, IL-2, MIP-1β) using the Boolean AND function.

The MHC association (MHC-Ia, MHC-E, MHC-II) of a 15mer peptide response was determined by pre-incubating isolated mononuclear cell aliquots for 1 hr at room temperature (prior to adding peptides or combining effector and target cells and incubating per the standard ICS assay) in the presence (and absence) of each the following specific inhibitors: 1) the pan anti-MHC-I mAb W6/32 (10μg/ml), 2) the MHC-II-blocking mAb G46.6 (10μg/ml), or 3) the MHC-E blocking VL9 peptide (VMAPRTLLL; 20μM). Stimulated cells were fixed, permeabilized, stained and analyzed as described above. To be considered MHC-E-restricted by blocking, the individual peptide response must have been blocked by both anti-pan MHC-I clone W6/32 and MHC-E-binding peptide VL9, and not blocked by anti-MHC-II. MHC-II-restricted responses were blocked by anti-MHC-II but not anti-MHC-I or VL9, and MHC-Ia-restricted responses were blocked by anti-MHC-Ia only (*9, 10*). Responses that did not meet these inhibition criteria were considered indeterminate. Minimal independent epitope numbers were estimated from the positive responses identified by testing of consecutive 15mer peptides by the following criteria: single positive peptide of same restriction type = 1 independent epitope; 2 adjacent positive peptides of same restriction type = 1 independent epitope; 3 adjacent positive peptides of same restriction type = 2 independent epitopes; 4 adjacent positive peptides of same restriction type = 2 independent epitopes; and 5 adjacent positive peptides of same restriction type = 3 independent epitopes.

**SIV detection assays.** Plasma SIV RNA levels were determined using an SIVgag-targeted quantitative real time/digital RT-PCR format assay, essentially as previously described, with 6 replicate reactions analyzed per extracted sample for assay thresholds of 15 SIV RNA copies/ml (*7, 8, 28*). Quantitative assessment of SIV DNA and RNA in cells and tissues was performed using SIVgag targeted, nested quantitative hybrid real-time/digital RT-PCR and PCR assays, as previously described (*7, 8, 28*). SIV RNA or DNA copy numbers were normalized based on quantitation of a single copy rhesus genomic DNA sequence from the CCR5 locus from the same specimen, as described, to allow normalization of SIV RNA or DNA copy numbers per $10^8$ diploid genome cell equivalents. Ten replicate reactions were performed with aliquots of extracted DNA or RNA from each sample, with two additional spiked internal control reactions performed with each sample to assess potential reaction inhibition. Samples that did not yield any positive results across the replicate reactions were reported as a value of "less than" the value that would apply for one positive reaction out of 10. Threshold sensitivities for individual specimens varied as a function of the number of cells or amount of tissue available and analyzed; for graphing consistency values are plotted with a common nominal sensitivity threshold.

Statistical Analysis. Boxplots show jittered points and a box from 1st to 3rd quartiles (IQR) and a line at the median, with whiskers extending to the farthest data point within 1.5*IQR above and below the box. For all comparisons of T cell response parameters, we performed Wilcoxon rank-sum tests comparing each non-RhCMV 68-1 vaccine group to the RhCMV 68.1 reference group. For longitudinal responses, we calculated the area under the curve (AUC) or the plateau average value for each RM, as denoted in the figure legends. All Wilcoxon P values are based on two-sided tests and unadjusted except where noted. Adjusted P values were computed using the Holm procedure for family-wise error rate control. All P values for analyses of efficacy or epitope restriction frequency were based on two-sided exact tests of binomial proportions. Analyses were performed in R v3.6.0 with the package Exact v2.0 (*29*).

**Supplemental Figures**

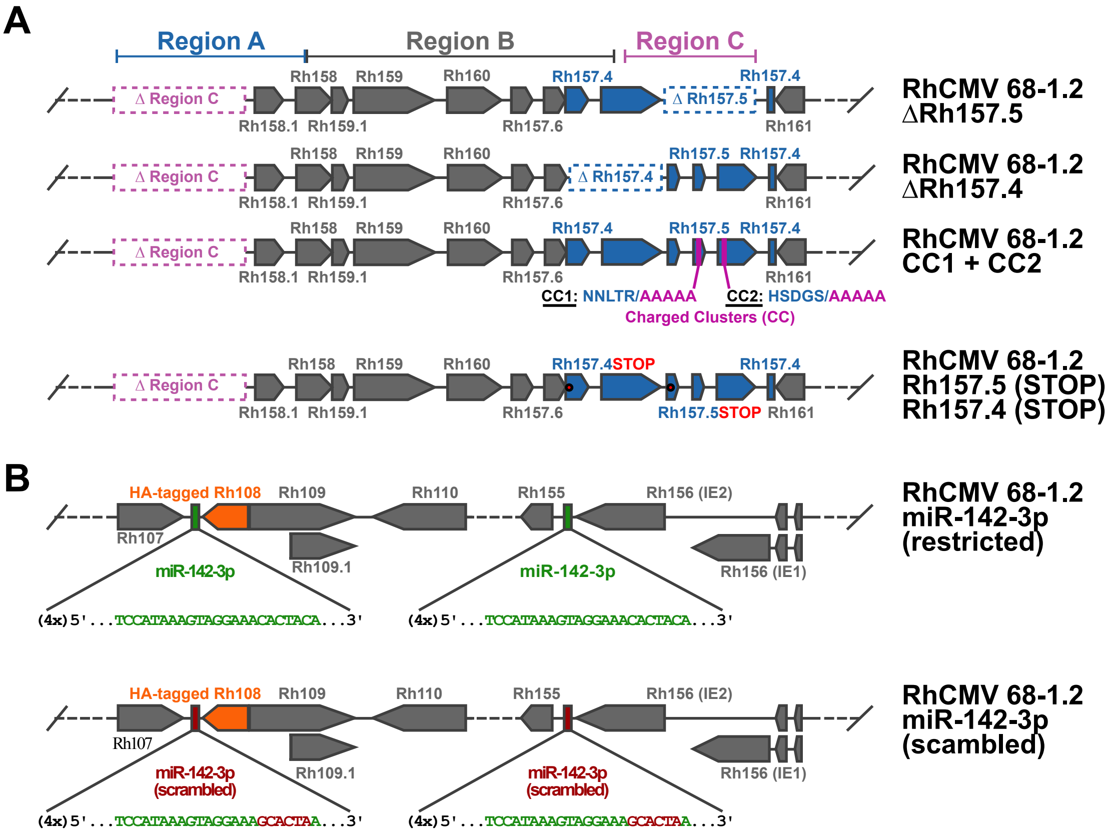


**Figure S1A-B. Genetic configuration of RhCMV vectors.** Schematic representation of the RhCMV genome modifications of the 68-1.2 RhCMV vectors used in this study. (**A**) Pictograms of genetic differences in the RhCMV genome region encoding the PRC (analogous to the $U_L$b' region of HCMV) of the modified RhCMV 68-1/68-1.2 vectors listed in **Fig. 1**. The genomic segments of inversion and deletion indicated by the letters A, B and C were adapted from Oxford *at al*. (*11*). Each pointed box represents a distinct exon, the point indicating the 3' end of the ORF. Highlighted boxes in dark gray, dark blue, and purple represent regions A, B, and C in the full length (wildtype) genetic configuration (**see Fig. 1A**). Dashed boxes represent functionally deleted viral ORFs or viral genomic segments indicated by (Δ). In RhCMV 68-1.2 157.5-CC1 + CC2 diagram, the changes in the charged clusters (CC1) NNLTR and (CC2) HSDGS located in the second and the third exons of the Rh157.5 gene, respectively, are indicated (replaced by AAAAA). Homologous charge clusters in UL128 of HCMV were previously shown to abolish PRC formation (14). Red dots in the last diagram represent inserted termination codons (STOP) in Rh157.5 and Rh157.4 of RhCMV 68-1.2. (**B**) Schematics highlight the genomic regions where intact or sequence-scrambled miR-142-3p recognition sites were inserted into RhCMV 68-1.2 for tropism restriction. Each green box represents an insertion of four miR-142-3p-restricted recognition sites (4x). Sequences highlighted in green represent the miR-142-3p-specific recognition site sequence. Red boxes indicate a replacement of the designated miR-142-3p target sequences highlighted in green with scrambled sequences highlighted in red. Orange boxes represent the Rh108 ORF with the addition of a hemagglutinin (HA) epitope-tag.

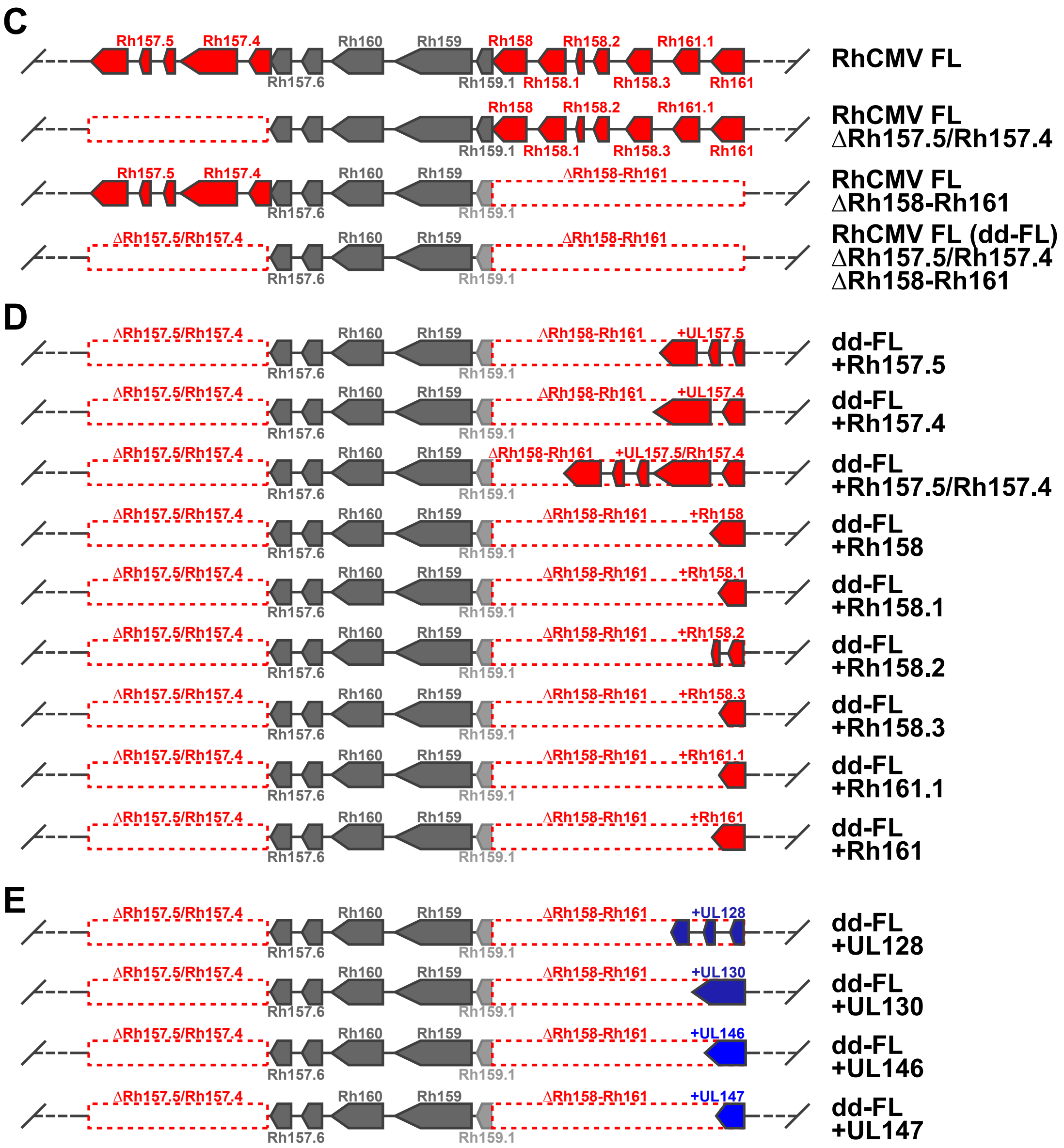


**Figure S1C-E. Genetic configuration of RhCMV vectors.** Schematic representation of the RhCMV genome modifications of the FL-RhCMV vectors used in this study. (**C-E**) Pictograms of the genetic differences in the RhCMV genome region encoding the PRC (analogous to the $U_L$b' region of HCMV) of the modified RhCMV FL vectors listed in **Fig. 2**. Each box represents a distinct exon and each dashed box highlighted in red represents a deleted region (Δ) of the corresponding genes highlighted in red in RhCMV FL. Double-deleted Full Length (**dd-FL**) is the designation for the RhCMV FL vector deleted for both Rh157.5/Rh157.4 and Rh158-Rh161 genomic regions.

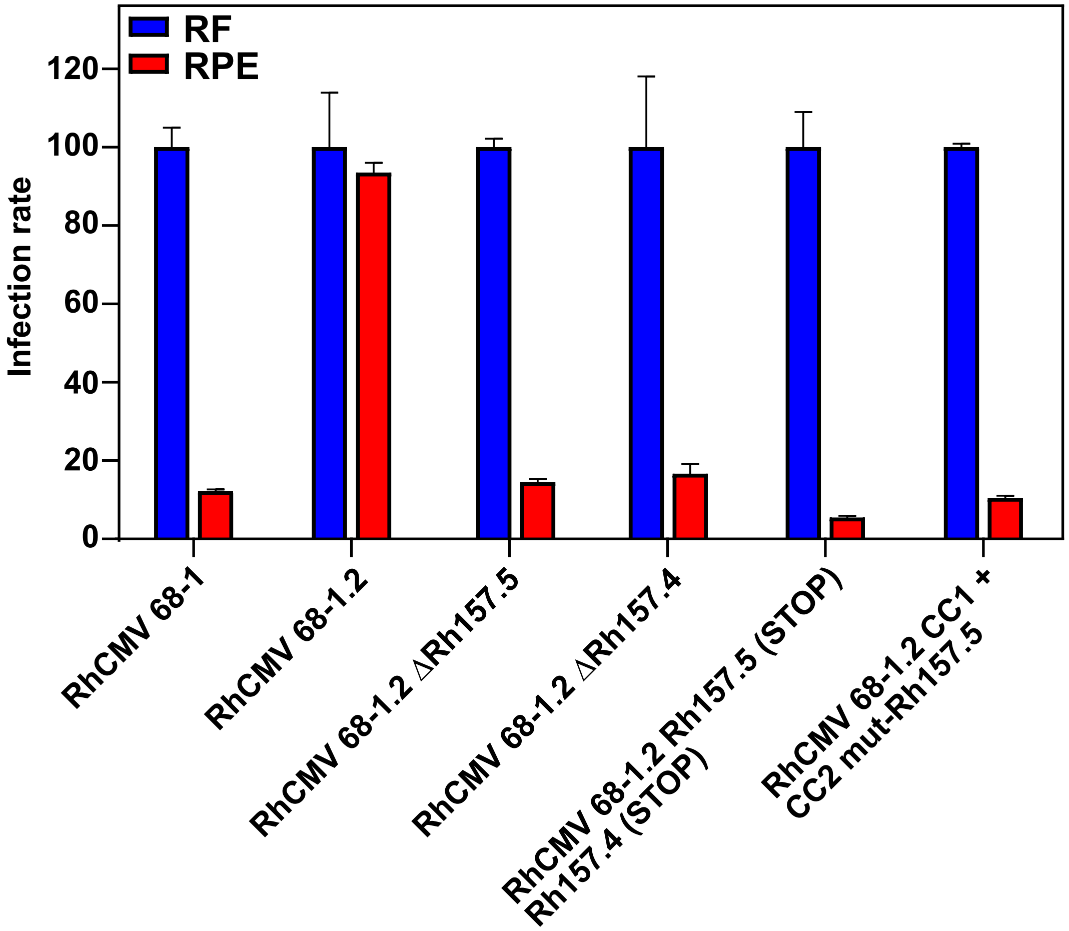


**Figure S2. Functional PRC analysis of 68-1.2-based RhCMV vectors.** The indicated vectors were used to infect triplicate repeat cultures of primary rhesus fibroblasts (RF) or rhesus retinal pigment epithelial cells (RPE) at multiplicities of infection of 0.3 and 10, respectively. All samples were harvested at 48 hours post infection and subsequently fixed and permeabilized before infection rates (% infected cells) were determined by flow cytometry using a RhCMV-specific antibody. Mean infection rates in RFs were set to 100% and infection rates in RPEs are shown in relation to RFs (+ SEM). Reduced infectivity of RPEs relative to RFs indicates abrogation of PRC function.

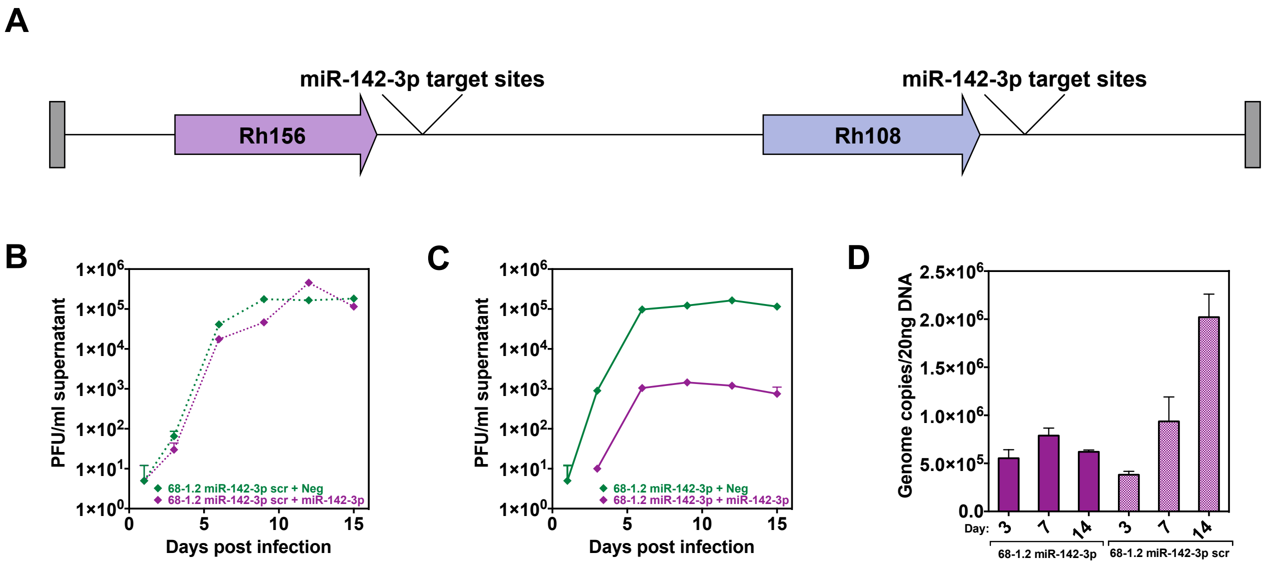


**Figure S3. Myeloid cell tropism-restriction of 68-1.2 RhCMV with microRNA (miR)-142-3p.** Characterization of the 68-1.2 miR-142-3p vaccine vector. (**A**) Schematic of the miR-142-3p target sites inserted 20 nucleotides downstream of the termination codons of both Rh156 and Rh108 encoding the conserved RhCMV homologs of HCMV essential genes IE2 and UL79. (**B**) Growth analysis of the 68-1.2 control vector containing a scrambled version of the miR-142-3p target sites in the presence of exogenous miR-142-3p. Primary rhesus fibroblasts were transfected with negative control or miR-142-3p mimic and then infected 24 hours later with 68-1.2 miR-142-3p scrambled control virus at MOI = 0.01. Cell supernatants were harvested at the indicated timepoints and titered on primary rhesus fibroblasts. (**C**) Growth analysis of the 68-1.2 miR-142-3p vector was performed as in panel B. (**D**) Analysis of RhCMV DNA replication in primary rhesus macrophages. Monocytes were derived from peripheral blood of 3 animals, differentiated into macrophages *in vitro* and infected with 68-1.2 miR-142-3p or scrambled virus at MOI = 5. At the indicated times post-infection viral DNA was isolated and total DNA copies were determined using qPCR.

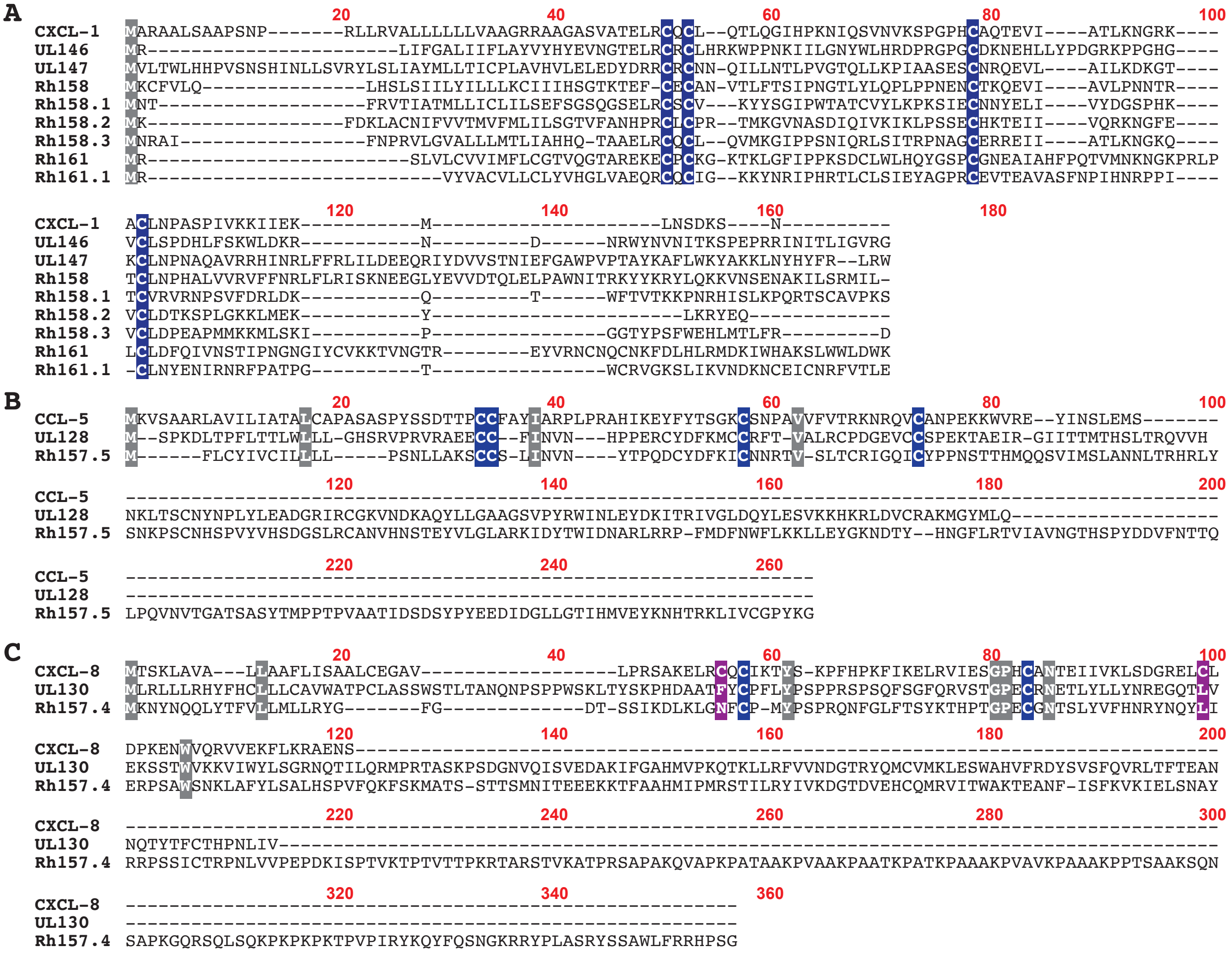


**Figure S4. Protein sequence comparison of RhCMV/HCMV orthologs RhCMV157.5/UL128, Rh157.4/ UL130 and Rh158-161/UL146-147.** Shown are complete protein sequence alignments of the HCMV/RhCMV encoded UL146 family members in comparison to human CXCL-1 (#NP_001502) (**A**), the UL128/Rh157.5 proteins in comparison to human CCL-5 (#NG_015990) (**B**) and the UL130/ Rh157.4 proteins in comparison to human CXCL-8 (#NG_029889) (**C**). The conserved structural cysteines important for chemokine function are highlighted in blue while all other conserved amino acids are highlighted in grey. Also highlighted in panel C (magenta) are two of the four cysteines characteristic for CXC-chemokines found in CXCL-8, but not in UL130/Rh157.4.

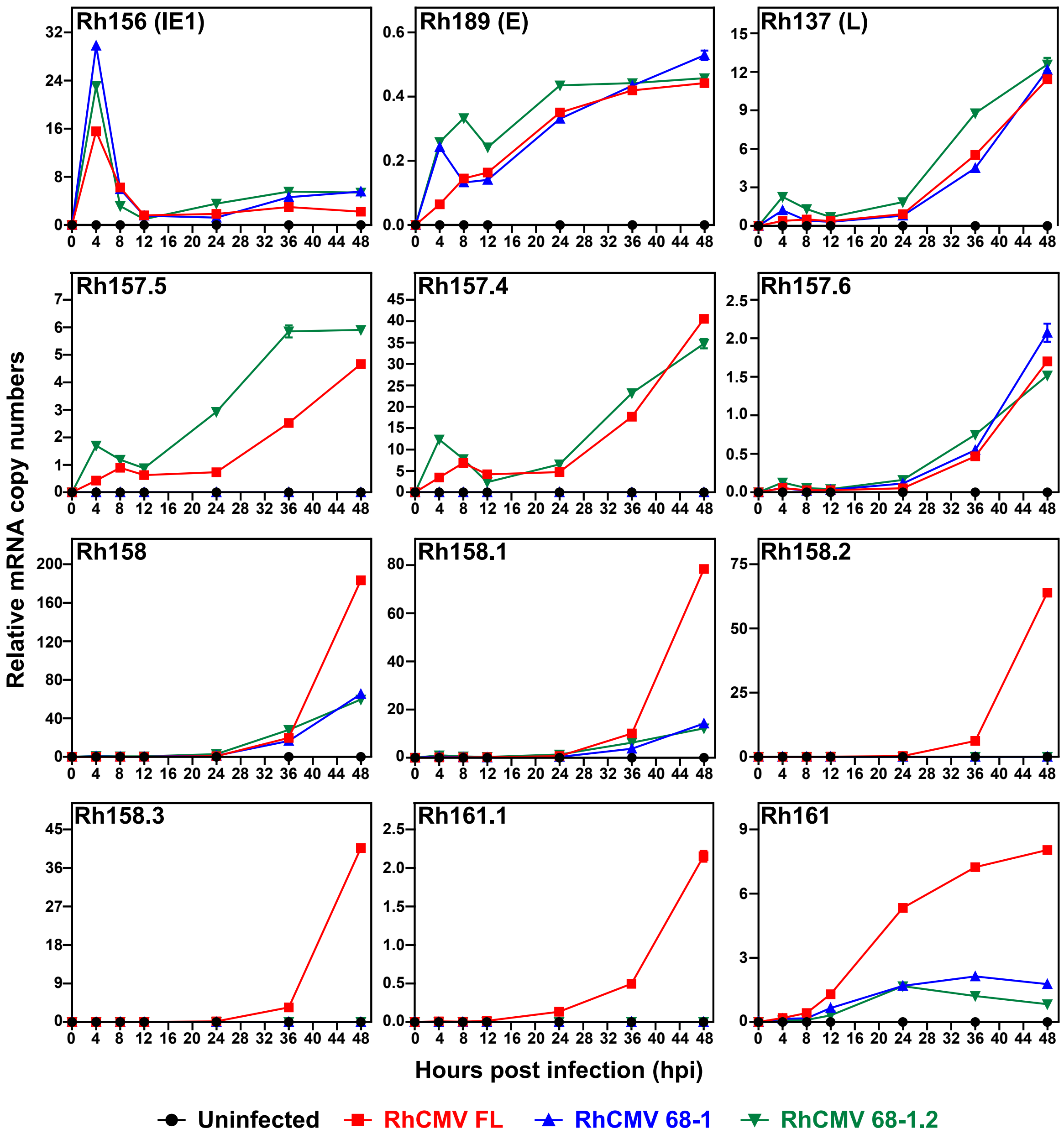


**Figure S5. Transcriptional analysis of Rh158-Rh161 region genes in RhCMV FL in comparison to RhCMV 68-1 and 68-1.2.** Total RNA was isolated from RM fibroblasts infected with a MOI = 5 of RhCMV-FL, RhCMV 68-1, or RhCMV 68-1.2 at 0, 4, 8, 12, 24, 36, and 48 hours post infection (hpi). Uninfected rhesus fibroblasts were used as a negative control. The mRNA copy number of each gene relative to the housekeeping gene GAPDH was determined by quantitative reverse transcriptase PCR (qRT-PCR) analysis at the indicated time points. As a positive control, the transcripts of Rh156, Rh189, and Rh137 (upper panels) representing immediate early (IE), early (E), and late (L) genes, respectively, were included. The lack of expression of the PRC genes Rh157.5 and Rh157.4 but not Rh157.6 was confirmed in RhCMV 68-1 infected cells whereas all three genes were co-expressed in 68-1.2 and RhCMV FL. Compared to RhCMV FL, quantification of the six Rh158-Rh161 region transcripts (lower two panels) confirmed the lack of Rh158.2, Rh158.3, and Rh161.1 gene expression in 68-1 and 68-1.2-infected cells and revealed substantially lower expression levels of Rh158, Rh158.1, and Rh161 genes at late times of infection.

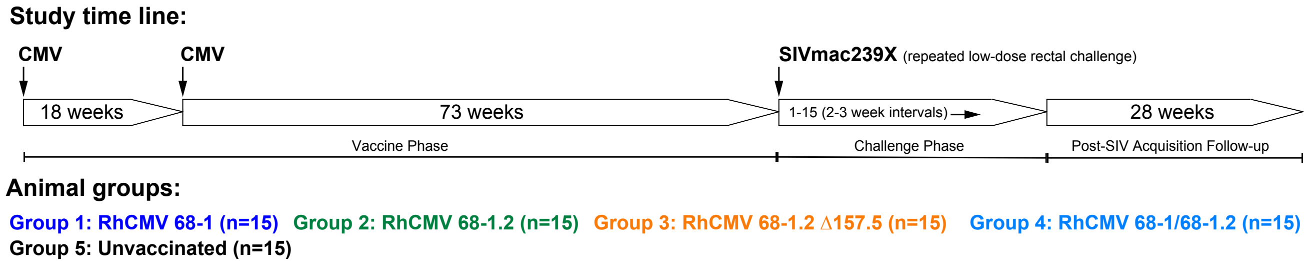


**Figure S6. Protocol for the comparison of the immunogenicity and efficacy of 68-1, 68-1.2, and ΔRh157.5 68-1.2 RhCMV/SIV vector sets, and the combination of 68-1 and 68-1.2 vector sets.** Study RM (n =15 per group) were subcutaneously vaccinated twice at the indicated timepoints with vector sets of the indicated RhCMV vector backbones. Each set was comprised of three RhCMV vectors of the same genetic backbone individually expressing SIVgag, rev/tat/nef fusion protein, or 5'-pol, administered at a dose of 5 x $10^6$ infectious units per vector**.** No dose adjustment was made for RM receiving both the 68-1 and 68-1.2 vector sets. Thus, these RM therefore received a total vaccine dose that was 2X that of RM receiving only one vector set. An additional n = 15 unvaccinated RM were included as negative controls. Repeated, intra-rectal, limiting dose SIVmac239 challenge was initiated on vaccinated RM 90 weeks after first vaccination and carried out for all groups until "take" of SIV infection as described in the *Methods*. After documentation of SIVmac239 "take", RM with progressive (continuously viremic) infection were followed for 10 weeks, whereas protected RM were followed 28 weeks.

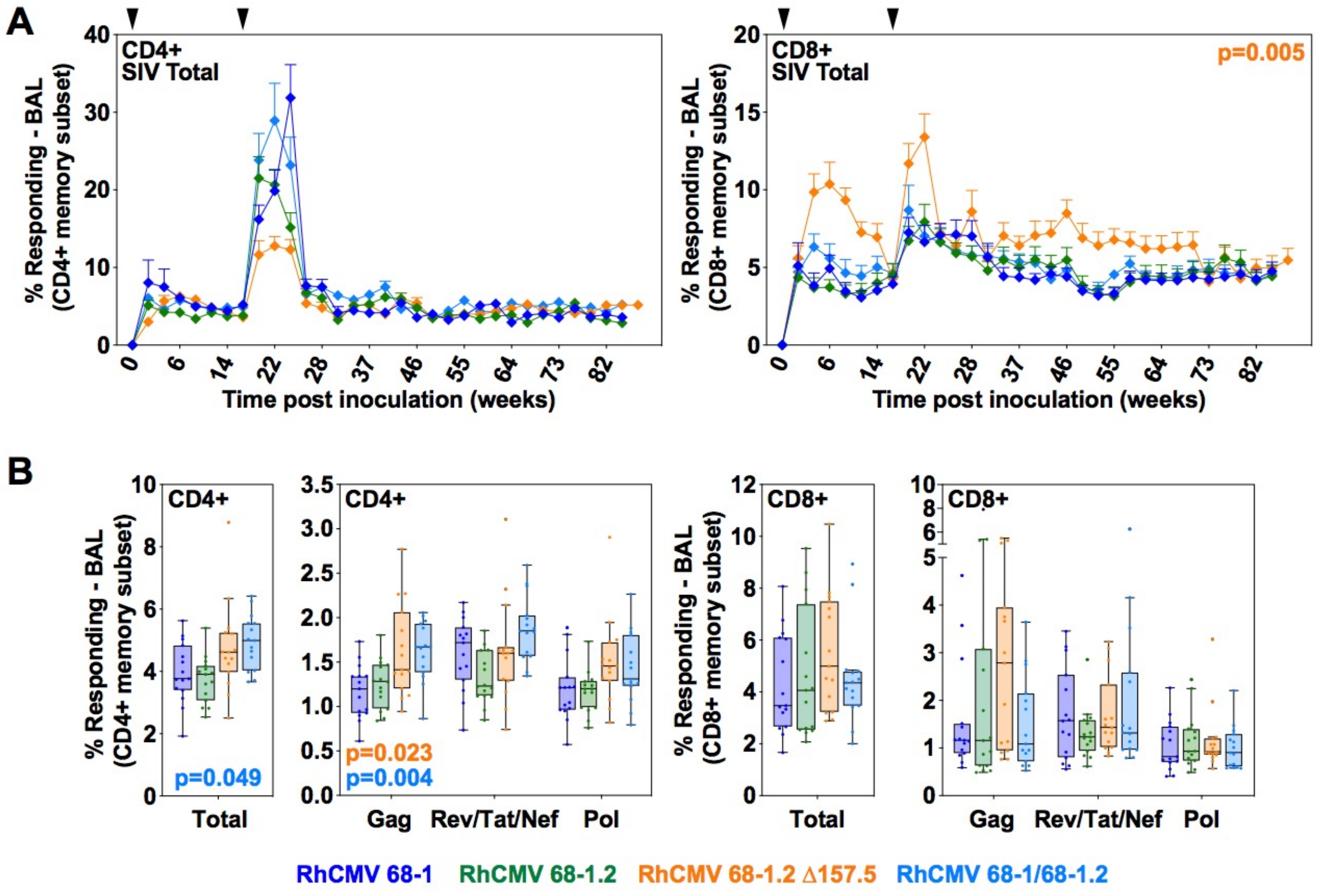


**Figure S7. Analysis of SIV-specific T cell responses in bronchoalveolar alveolar lavage fluid (BAL) in RM cohorts destined for SIV challenge.** (**A**) Longitudinal analysis of the total vaccine-elicited SIV -specific CD4+ and CD8+ T cell responses (sum of Gag, Rev/Tat/Nef, and 5'-Pol-specifc responses) in BAL of the RM vaccinated with 68-1, 68-1.2, and 68-1.2 ΔRh157.5 vector sets and the combination of the 68-1 and 68-1.2 vector sets. Responses to whole SIV protein peptide mixes were assessed by a flow cytometric ICS assay (based on induction of TNF and/or IFN-γ) as described in **Fig. 3**. (**B**) Plateau phase analysis of the same ICS data showing total SIV-specific responses and responses to each individual SIV insert. Each data point is the mean of response frequencies in all samples from 61-88 weeks post-first vaccination. Wilcoxon p-values for comparison of the 68-1-only vaccine to the other vaccines are shown where significant (unadjusted in Panel A, adjusted across 3 insert Ags in Panel B).

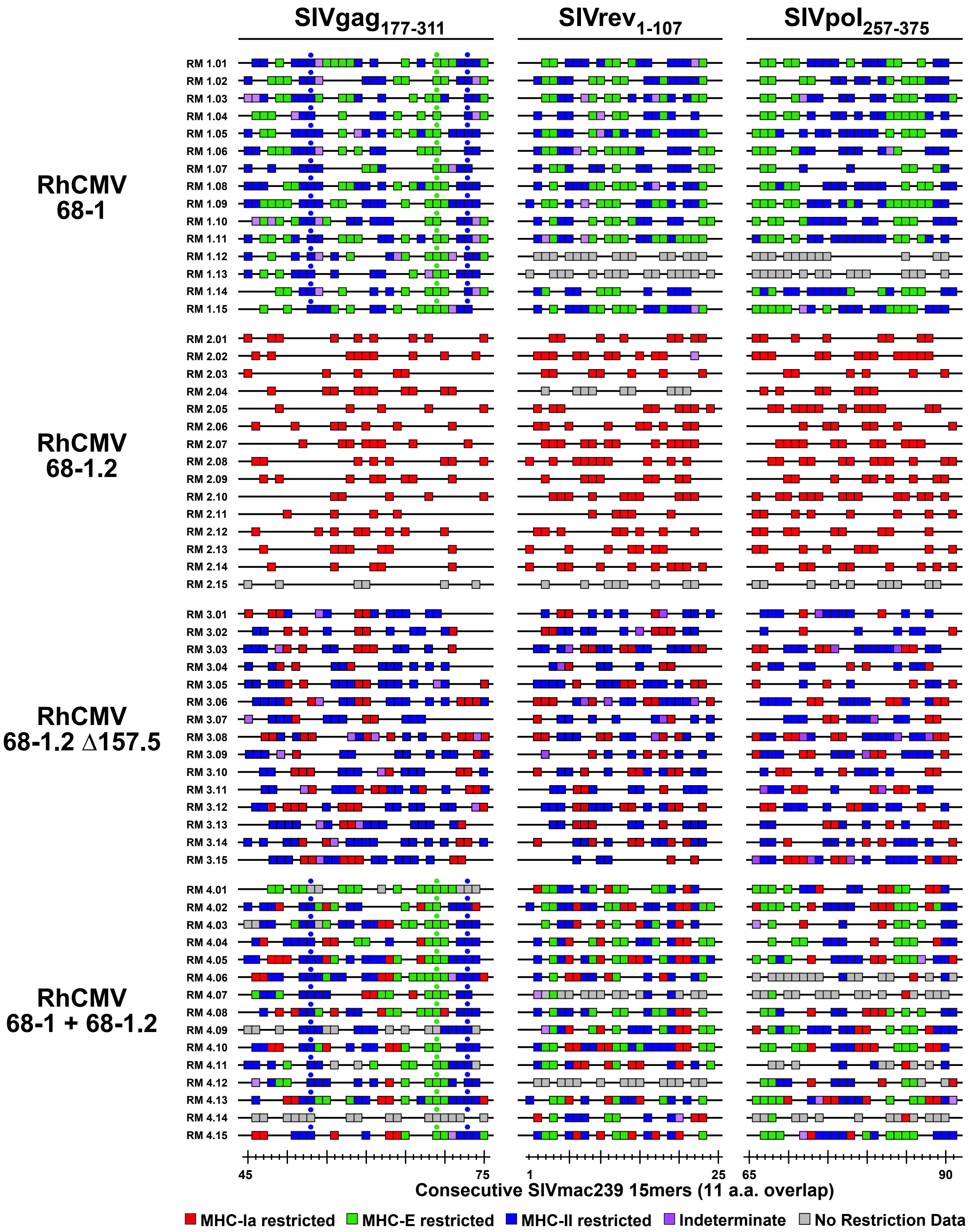


**Figure S8. Epitope analysis of RhCMV vaccine-elicited, SIV-specific CD8+ T cell responses in RM cohorts destined for SIV challenge.** The epitope targeting characteristics of the CD8+ T cell responses elicited by each vector set (RhCMV/SIVgag, RhCMV/SIVretanef, and RhCMV/SIV5'pol) in the RM cohorts destined for SIV challenge were analyzed for indicated portions of each insert ($SIVgag_{177-311}$, $SIVrev1_{-107}$, $SIVpol_{257-375}$ for the SIVgag, SIVretanef, and SIV5'-pol inserts, respectively, corresponding to 15mers 45-75, 1-24, and 65-91). Grey boxes reflect insufficient cell availability to complete restriction analysis. Blue and green dots indicate independently confirmed MHC-II- and MHC-E-restricted supertope responses, respectively.

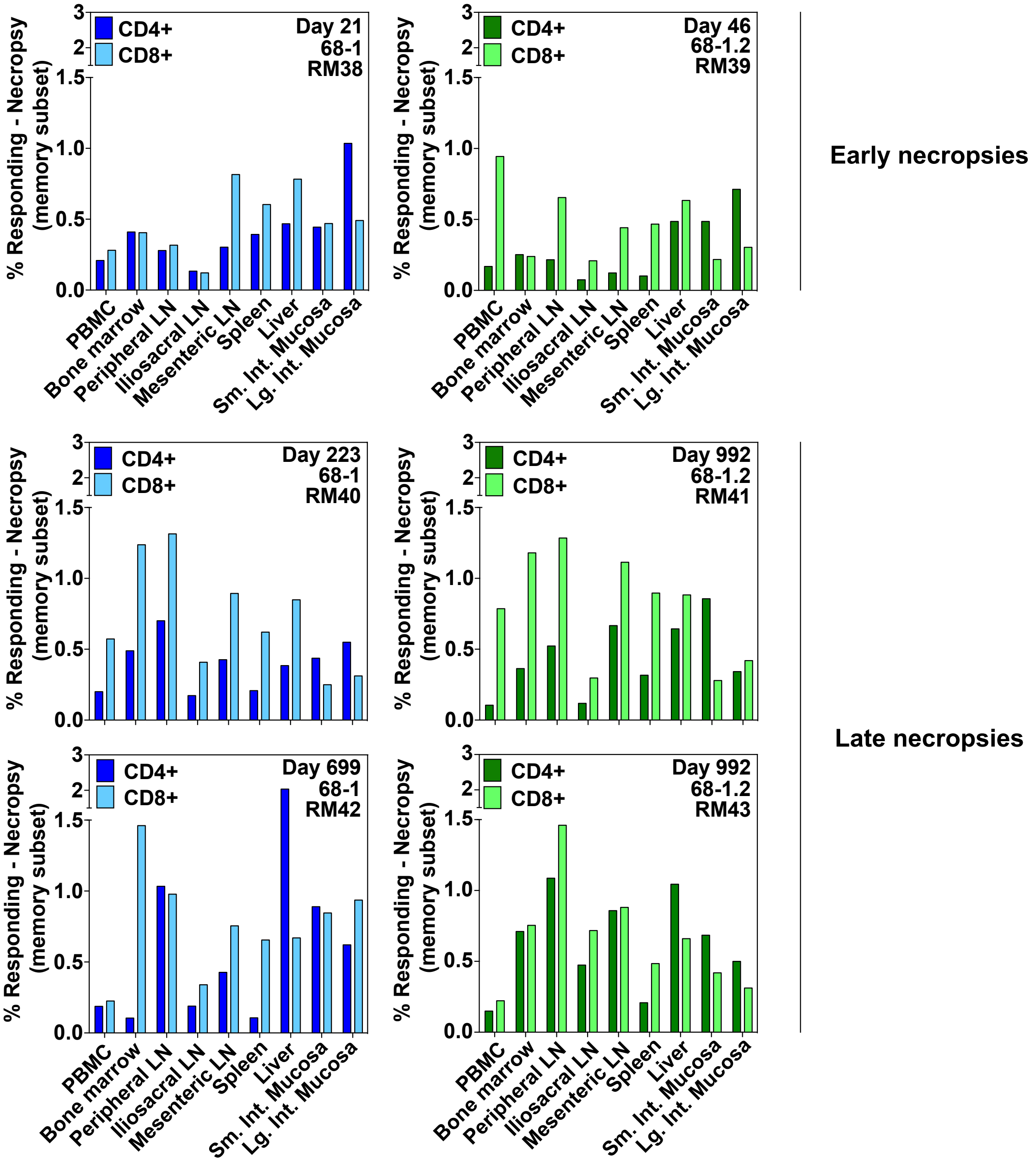


**Figure S9. Analysis of SIVgag-specific CD4+ and CD8+ T cell response magnitude in tissues of 68-1 vs. 68-1.2 RhCMV/SIVgag vector vaccinated RM at necropsy.** Mononuclear cells from blood and the indicated tissues from RM inoculated with either the 68-1 or 68-1.2 RhCMV/SIVgag vectors were obtained at necropsy for analysis of SIV Gag-specific CD4+ and CD8+ T cell responses. Responses to whole SIV Gag protein peptide mix were assessed by a flow cytometric ICS assay (based on induction of TNF and/or IFN-γ) as described in **Fig. 3**.

| Vector | # RM | assigned peptide responses | restriction-assigned epitopes | # MHC-Ia-restricted epitope responses (%) | # MHC-E-restricted epitope responses (%) | # MHC-II-restricted epitope responses (%) |
|---|---|---|---|---|---|---|
| 68-1 | 15 | 1047 | 704 | 0 (0) | 320 (45.5) | 384 (54.5) |
| 68-1.2 | 14 | 613 | 418 | 418 (100) | 0 (0) | 0 (0) |
| 68-1.2 Rh157.5 (STOP) Rh157.4 (STOP) | 2 | 115 | 78 | 0 (0) | 39 (50.0) | 39 (50.0) |
| 68-1.2 ΔRh157.5 | 15 | 391 | 267 | 104 (39.0) | 0 (0) | 163 (61.0) |
| 68-1.2 ΔRh157.4 | 4 | 219 | 157 | 72 (45.9) | 0 (0) | 85 (54.1) |
| 68-1.2 CC1 + CC2 mut-Rh157.5 | 2 | 105 | 70 | 34 (48.6) | 0 (0) | 36 (51.4) |
| 68-1.2 miR-142-3p-restricted | 3 | 143 | 95 | 11 (11.6) | 0 (0) | 84 (88.4) |
| 68-1.2 miR-control (142-3p scrambled) | 2 | 103 | 73 | 73 (100) | 0 (0) | 0 (0) |
| 68-1 + 68-1.2 | 14 | 940 | 641 | 136 (21.2) | 220 (34.3) | 285 (44.5) |
| RhCMV FL | 2 | 120 | 76 | 76 (100) | 0 (0) | 0 (0) |
| FL ΔRh157.5/Rh157.4 | 3 | 167 | 108 | 108 (100) | 0 (0) | 0 (0) |
| FL ΔRh158-Rh161 | 2 | 98 | 68 | 68 (100) | 0 (0) | 0 (0) |
| dd FL* | 3 | 192 | 124 | 0 (0) | 60 (48.4) | 64 (51.6) |
| dd FL + Rh157.5 | 1 | 55 | 37 | 17 (45.9) | 0 (0) | 20 (54.1) |
| dd FL + Rh157.4 | 1 | 61 | 40 | 23 (57.5) | 0 (0) | 17 (42.5) |
| dd FL + Rh157.5/Rh157.4 | 2 | 111 | 70 | 70 (100) | 0 (0) | 0 (0) |
| dd FL + Rh158 | 1 | 53 | 34 | 34 (100) | 0 (0) | 0 (0) |
| dd FL + Rh158.1 | 1 | 49 | 35 | 35 (100) | 0 (0) | 0 (0) |
| dd FL + Rh158.2 | 1 | 55 | 35 | 35 (100) | 0 (0) | 0 (0) |
| dd FL + Rh158.3 | 1 | 53 | 35 | 35 (100) | 0 (0) | 0 (0) |
| dd FL + Rh161.1 | 1 | 55 | 37 | 37 (100) | 0 (0) | 0 (0) |
| dd FL + Rh161 | 1 | 62 | 41 | 41 (100) | 0 (0) | 0 (0) |
| dd FL + HCMV UL128 | 2 | 101 | 74 | 35 (47.3) | 0 (0) | 39 (52.7) |
| dd FL + HCMV UL130 | 2 | 99 | 73 | 34 (46.6) | 0 (0) | 39 (53.4) |
| dd FL + HCMV UL146 | 2 | 98 | 72 | 72 (100) | 0 (0) | 0 (0) |
| dd FL + HCMV UL147 | 2 | 114 | 78 | 78 (100) | 0 (0) | 0 (0) |
| dd FL + HCMV UL146 STOP | 2 | 112 | 79 | 0 (0) | 38 (48.1) | 41 (51.9) |

*RhCMV FL ΔRh157.5/Rh157.4 + ΔRh158-Rh161

**Table S1. Epitope analysis summary of all study RM.** The table shows the total numbers of SIV Gag 15mer peptide-specific $CD8^+$ T cell responses that were restriction-assignable for each of the designated RhCMV/SIVgag vectors, the corresponding number of independent epitopes, and the restriction type of each of the independent epitope-specific responses (see Methods).
